# Bacterial vaginosis-driven changes in cervicovaginal immunity that expand the immunological hypothesis for increased HIV susceptibility

**DOI:** 10.1101/2024.07.03.601916

**Authors:** Finn MacLean, Adino Tesfahun Tsegaye, Jessica B. Graham, Jessica L. Swarts, Sarah C. Vick, Nicole Potchen, Irene Cruz Talavera, Lakshmi Warrier, Julien Dubrulle, Lena K. Schroeder, Ayumi Saito, Katherine K. Thomas, Matthias Mack, Michelle C. Sabo, Bhavna H. Chohan, Kenneth Ngure, Nelly Mugo, Jairam R. Lingappa, Jennifer M. Lund, the Kinga Study Team

**Affiliations:** Vaccine and Infectious Disease Division, Fred Hutchinson Cancer Center, Seattle, USA; Department of Global Health, University of Washington, Seattle, USA; Cellular Imaging Shared Resource, Fred Hutchinson Cancer Research Center, Seattle, USA; Department of Internal Medicine-Nephrology, University Hospital Regensburg, Regensburg, Germany; Department of Medicine, University of Washington, Seattle, USA; Center for Virus Research, Kenya Medical Research Institute, Nairobi, Kenya; School of Public Health, Jomo Kenyatta University of Agriculture and Technology, Nairobi, Kenya; Center for Clinical Research, Kenya Medical Research Institute, Nairobi, Kenya; Department of Pediatrics, University of Washington, Seattle, USA

## Abstract

Bacterial vaginosis (BV) is a dysbiosis of the vaginal microbiome that is prevalent among reproductive-age females worldwide. Adverse health outcomes associated with BV include an increased risk of sexually-acquired HIV, yet the immunological mechanisms underlying this association are not well understood. To investigate BV-driven changes to cervicovaginal tract (CVT) and circulating T cell phenotypes, participants with or without BV provided vaginal tract (VT) and ectocervical (CX) tissue biopsies and PBMC samples. High-parameter flow cytometry revealed an increased frequency of cervical conventional CD4^+^ T cells (Tconv) expressing CCR5. However, we found no difference in number of CD3^+^CD4^+^CCR5^+^ cells in the CX or VT of BV+ vs BV- individuals, suggesting that BV-driven increased HIV susceptibility may not be solely attributed to increased CVT HIV target cell abundance. Flow cytometry also revealed that individuals with BV have an increased frequency of dysfunctional CX and VT CD39^+^ Tconv and CX tissue-resident CD69^+^CD103^+^ Tconv, reported to be implicated in HIV acquisition risk and replication. Many soluble immune factor differences in the CVT further support that BV elicits diverse and complex CVT immune alterations. Our comprehensive analysis expands on potential immunological mechanisms that may underlie the adverse health outcomes associated with BV including increased HIV susceptibility.

## Introduction

Bacterial vaginosis (BV) is characterized by a vaginal dysbiosis in which the normally *Lactobacillus*-dominated microbiome is replaced by one with higher bacterial diversity, including increased concentrations of anaerobic bacteria (1). BV results in symptoms of vaginal inflammation, discharge, and discomfort, and is estimated to affect 23-29% of reproductive-age females worldwide (2), making BV the most common cause of vaginal symptoms leading patients to seek medical care (3). BV is also associated with several adverse medical outcomes (1, 2, 4-11) including preterm delivery (12), chorioamnionitis (13), endometritis (14), and an increased risk of HIV acquisition (5, 11). The issue of HIV acquisition risk is of particular concern in parts of the world where both BV and HIV are prevalent, such as in sub-Saharan Africa (15). Although many studies have investigated immune system alterations associated with BV (1, 5-7, 10, 11, 16-19), the link between CVT T cells, which may mediate the relationships between BV and adverse health outcomes, is not well understood. Here, we sought to better understand the impact of BV on CVT mucosal and systemic immune responses in Kenyans and how these changes may alter HIV susceptibility.

In the CVT, innate and adaptive immune cells line the epithelium and lamina propria to provide immunological surveillance (20). Notably, T cell populations in the CVT maintain distinct phenotypes from those circulating in the blood (21, 22). These include tissue-resident memory T cells (T_RM_) bearing CD69 with or without CD103 (23). CD69 is a marker of activation (24) that has a unique role in maintaining tissue residency by binding S1PR1 and preventing egress from the tissue (25-27). CD103 further promotes tissue residency by binding to E-cadherin (28). T_RM_ are retained in the mucosal tissue and have the ability to readily respond to pathogens *in situ* (21, 29-31). Upon recognition of their cognate antigen, T_RM_ use their “sensing and alarm function” to produce pro-inflammatory cytokines and chemokines that rapidly recruit and activate other immune cells, thereby potentiating the tissue immune response while also executing their specific effector and cytotoxic functions (32-35).

Given that in the context of heterosexual transmission, genital mucosal tissue is where host exposure to HIV most commonly first occurs, activated CD4^+^ T cells expressing the HIV coreceptor CCR5 in the CVT have been thought to be prime targets for initial HIV infection (36). Studies using a mouse model demonstrated that the introduction of BV-associated bacteria into the vagina led to an increased quantity of CVT CD4^+^ T cells expressing the activation marker CD44 and HIV coreceptor CCR5 (7), suggesting dysbiosis of normal vaginal flora may lead to an increase of HIV target cells. In humans, studies of immune responses in the context of BV have characterized immune cells collected from the CVT lumen via minimally invasive sampling techniques such as cervicovaginal lavage, vaginal swabs, cervical swabs, or cytobrushes. These studies have identified an increase in CCR5^+^CD4^+^ T cells in the CVT lumen of BV+ individuals (37-40), without addressing whether this observation is true in the CVT deeper tissue layers. In addition to the detection of HIV target cells in the CVT lumen of individuals with BV, studies have shown a connection between BV and the presence of pro-inflammatory cytokines in the CVT, most reliably IL-1β (7, 37, 41, 42); adding further support to the idea that proinflammatory mechanisms underlie the link between BV and increased HIV susceptibility. However, the relationship between BV and an increase in CD4^+^ T cell activation is inconsistently observed (43, 44), suggesting more complex mechanisms may be contributing to increased HIV susceptibility in individuals with BV. Further characterizing BV-driven immune modulations in the unique immune environment of the CVT is paramount to understanding mechanisms underlying increased HIV susceptibility in individuals with BV.

To comprehensively evaluate immune alterations associated with BV, we used a recent advance in tissue cryopreservation (45) combined with high-parameter, high-throughput flow cytometry to compare immune cells isolated from CVT tissue biopsies and blood from individuals with versus without BV. Given the dense T cell populations within CVT tissue (46), cells isolated from CVT mucosal biopsies may better represent the mucosal tissue immune environment than those isolated from the CVT lumen by cytobrush or other methods. Additionally, improved flow cytometry technology allows us to broadly evaluate immune cells in the context of BV for phenotypic markers not previously characterized.

Here, we report detailed T cell characteristics of vaginal tract (VT) and ectocervix (CX) mucosal tissues and PBMC samples from the same participants, comparing individuals with to those without BV. Additionally, we analyzed CVT fluids and serum for local and systemic cytokines and chemokines in individuals with versus without BV. While we did not observe an overall increase in CD3^+^CD4^+^CCR5^+^ HIV target cells in the CVT of BV+ individuals, we did observe CX and VT T cells with dysfunctional phenotypes and altered expression of soluble mediators in CVT fluid from BV+ individuals. Our findings highlight additional novel mechanisms that may underlie increased HIV susceptibility associated with BV.

## Results

### Study population characteristics

204 participants met the criteria to be included in the BV flow cytometry and/or soluble immune factor analysis (**Table 1**). 46 participants contributed at least one sample when they were BV+ by a Nugent score of 7-10 (34 at study enrollment and 12 at study exit). The BV- comparator group comprised N=158 participants with normal flora (Nugent score 0-3) who provided at least one sample at enrollment. Overall, our study population was young (55.9% were <30 years of age), sexually active (median of 8 unprotected sex acts in the prior month), with 40.8% using hormonal contraceptives, 39.9% Herpes Simplex Virus 2 (HSV-2) seropositive, and 17.6% had a sexual partner living with HIV (**Table 1**). 187 participants provided a CX sample (N=149 BV-, N=38 BV+), 190 participants provided a VT sample (N=149 BV-, N=41 BV+), and 185 participants provided a PBMC sample (N=143 BV-, N=42 BV+) for flow cytometry analysis. 193 participants provided a serum sample (N=149 BV-, and N=44 BV+) and 174 participants provided a CVT fluid sample (N= 136 BV-, N=38 BV+) for soluble immune factor analysis. A subset of individuals who were HSV-2 seronegative and HIV unexposed individuals provided a VT and/or CX tissue sample for immunofluorescent imaging at their enrollment visit (N=55 VT BV-, N=12 VT BV+, N=44 CX BV-, N=11 CX BV+).

**Table 1.** Demographic and Clinical Characteristics. Demographic and clinical data from the 204 participants who were included in the flow cytometry and cytokine analyses.

| **BV Status** | **Overall**<br>**N = 204<sup>1</sup>** | **Negative**<br>**N = 158<sup>1</sup>** | **Positive**<br>**N = 46<sup>1</sup>** |
| --- | --- | --- | --- |
| **Age** |  |  |  |
| <=24 | 66 (32.4%) | 48 (30.4%) | 18 (39.1%) |
| 25-29 | 48 (23.5%) | 36 (22.8%) | 12 (26.1%) |
| 30-34 | 33 (16.2%) | 27 (17.1%) | 6 (13.0%) |
| 35-39 | 29 (14.2%) | 23 (14.6%) | 6 (13.0%) |
| 40-49 | 27 (13.2%) | 23 (14.6%) | 4 (8.7%) |
| >=50 | 1 (0.5%) | 1 (0.6%) | 0 (0%) |
| **HIV Exposure** |  |  |  |
| HIV exposed (HIV serodifferent couple) | 36 (17.6%) | 23 (14.8%) | 13 (28.3%) |
| HIV unexposed (HIV negative, concordant couple) | 168 (82.4%) | 135 (85.4%) | 33 (71.7%) |
| **HSV-2 Status** |  |  |  |
| Seropositive | 81 (39.9%) | 56 (35.7%) | 25 (54.3%) |
| Seronegative | 114 (56.2%) | 95 (60.5%) | 19 (41.3%) |
| Indeterminate | 8 (3.9%) | 6 (3.8%) | 2 (4.3%) |
| Not Done | 1 | 1 | 0 |
| **Hormonal Contraceptive Use** |  |  |  |
| Had menses in last 35 days | 90 (44.8%) | 64 (40.8%) | 26 (59.1%) |
| Secondary amenorrhea | 29 (14.4%) | 24 (15.3%) | 5 (11.4%) |
| Uses hormonal contraceptives | 82 (40.8%) | 69 (43.9%) | 13 (29.5%) |
| Missing | 3 | 1 | 2 |
| **Number of unprotected vaginal intercourse**<br>**in last 30 days** | 8 (3,12) | 8 (3,12) | 8 (1,12) |
| <sup>1</sup>n (%); Median (IQR) |  |  |  |

### BV did not alter the balance of T cell frequency in the cervicovaginal tract tissues

To investigate the CVT mucosal tissue and circulating T cell population in detail, we performed high-parameter flow cytometry on cells isolated from cryopreserved CX and VT tissue biopsies cryopreserved using a previously published method (45), as well as PBMC samples. PBMC comparisons were *a priori* adjusted for hormonal contraceptive use, and CX and VT comparisons were *a priori* adjusted for hormonal contraceptive use, HSV-2 serology, HIV exposure, and semen exposure to reduce the effects of potential confounding variables on the analysis of BV-driven T cell alterations. We found that the majority of CD45^+^ cells in the VT and CX are CD3^+^ T cells (median > 75%; **Figure 1A**, **Supplemental Figure 1, Supplemental Table 1),** demonstrating the importance of T cells in immunosurveillance of the CVT. We first assessed whether BV was associated with a shift in the proportion of CD25^-^CD127^+/-^ conventional CD4^+^ T cells (Tconv) as a fraction of CD45^+^ cells in all three specimen types and found no significant difference comparing BV+ vs BV- samples (**Figure 1B**). The proportion of CD8^+^ T cells was analyzed as a fraction of CD45^+^ cells (**Figure 1C**), and no significant differences were detected comparing each sample type from BV+ vs BV- individuals. The fraction of Tconv cells of total CD3^+^ T cells (**Figure 1D**) and CD8^+^ of total CD3^+^ T cells (**Figure 1E**) were also not significantly altered by BV for any of the three sample types analyzed, indicating that BV may not be associated with a shift in the major T cell subsets.

**Figure 1.**
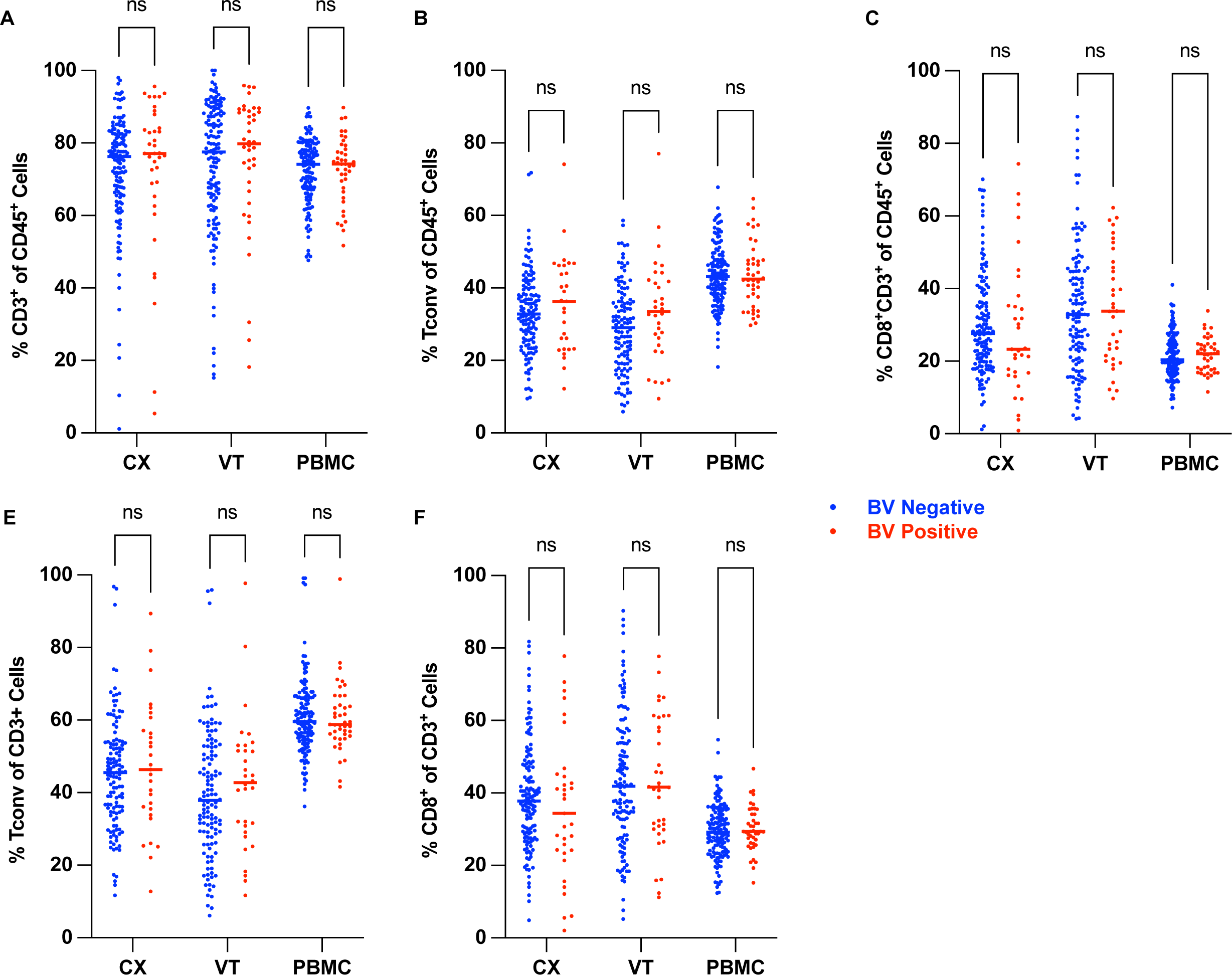
BV did not alter the balance of T cell frequency in the cervicovaginal tract tissues. Flow cytometry was used to quantify the proportions of T cells within different tissue sites as indicated. (**A**) The frequency of CD3^+^ among the total CD45^+^ population in ectocervix (CX), vaginal tract (VT), and peripheral mononuclear blood cells (PBMC) samples in BV- and BV+ individuals. The frequency of Tconv (CD3^+^CD4^+^CD25^-^ cells) (**B**) and CD3^+^CD8^+^ (**C**) among the total CD45^+^ population in CX, VT, and PBMC samples in BV- and BV+ individuals. The frequency of Tconv (**D**) and CD8^+^ (**E**) among the total CD3^+^ population in CX, VT, and PBMC samples in BV- and BV+ individuals. Adjusted rank regression analysis was performed to compare frequencies in each tissue between BV- and BV+ individuals. PBMC comparisons were *a priori* adjusted for hormonal contraceptive use, and CX and VT comparisons were *a priori* adjusted for hormonal contraceptive use, HSV-2 serology, HIV exposure, and semen exposure to reduce the effects of potential confounding variables on the analysis of BV-driven T cell alterations. Adjusted p-value displayed in bold when p ≤ 0.05, non-bold when 0.05 **<** p_adj_ ≤ 0.10, and “ns” for not significant when adjusted p **>** 0.10. Each dot represents a measurement from an individual sample. Each horizontal bar indicates the median for its respective group. For each comparison, the Ns, medians, p-values (adjusted and unadjusted), and estimated differences (adjusted and unadjusted) are provided in Supplemental Table 1.

### Conventional CD4^+^ T cells displayed increased markers of activation and tissue residency in the cervix of individuals with BV

We hypothesized that BV may be associated with phenotypic alterations among T cell subsets to promote adverse health outcomes. Previous studies have limited the characterization of cervicovaginal T cells to markers of inflammation, including the HIV co-receptor CCR5. Herein, we include markers of activation previously described to be enriched on CD4^+^ T cells of the cervicovaginal tract in the context of persistent BV, including CCR5, HLA-DR, and CD38 (44). However, we additionally sought to broaden our characterization of T cell phenotypic alterations associated with BV. We included Treg lineage markers (CD25, CD127, FoxP3) (47), a Th1 lineage marker (T-bet) (48), Th17 lineage markers (CD161, CCR6) (49, 50), markers of tissue residency (CD69 and CD103) (23), markers of inhibition (CD101 and CTLA-4) (51, 52), dysfunction (CD39) (53), progenitor potential (TCF-1) (54), exhaustion (PD-1) (55), an additional chemokine receptor used for trafficking and associated with activation (CXCR3) (56), cytotoxic function (Granzyme B) (57), and memory subsets (CCR7 and CD45RA) (58) to more broadly evaluate the phenotypic differences of CVT and circulating T cells in those with versus those without BV.

First, we evaluated whether BV is associated with activated Tconv phenotypes that may contribute to increased HIV susceptibility through a previously hypothesized mechanism – CCR5 expression. We found that CX CD4^+^ Tconv cells displayed a higher frequency of the HIV coreceptor CCR5 in BV+ individuals (median frequency 59% BV- vs 75% BV+, p_adj_ = 0.021; **Supplemental Table 1)** and minimal differences in the VT or PBMC samples as detected by flow cytometry and calculated as a fraction of total Tconv cells (**Figure 2A**). We also observed a significantly increased frequency of the activation marker HLA-DR on CX Tconv (median frequency 22% BV- vs 30% BV+, p_adj_ = 0.012; **Supplemental Table 1**), with minimal differences on the VT and PBMC Tconv cells in BV+ individuals (**Figure 2B**). Together, these results demonstrate that BV does promote local proinflammatory Tconv activation and that Tconv activation is elevated in the CX mucosa but not in the VT or circulation.

**Figure 2.**
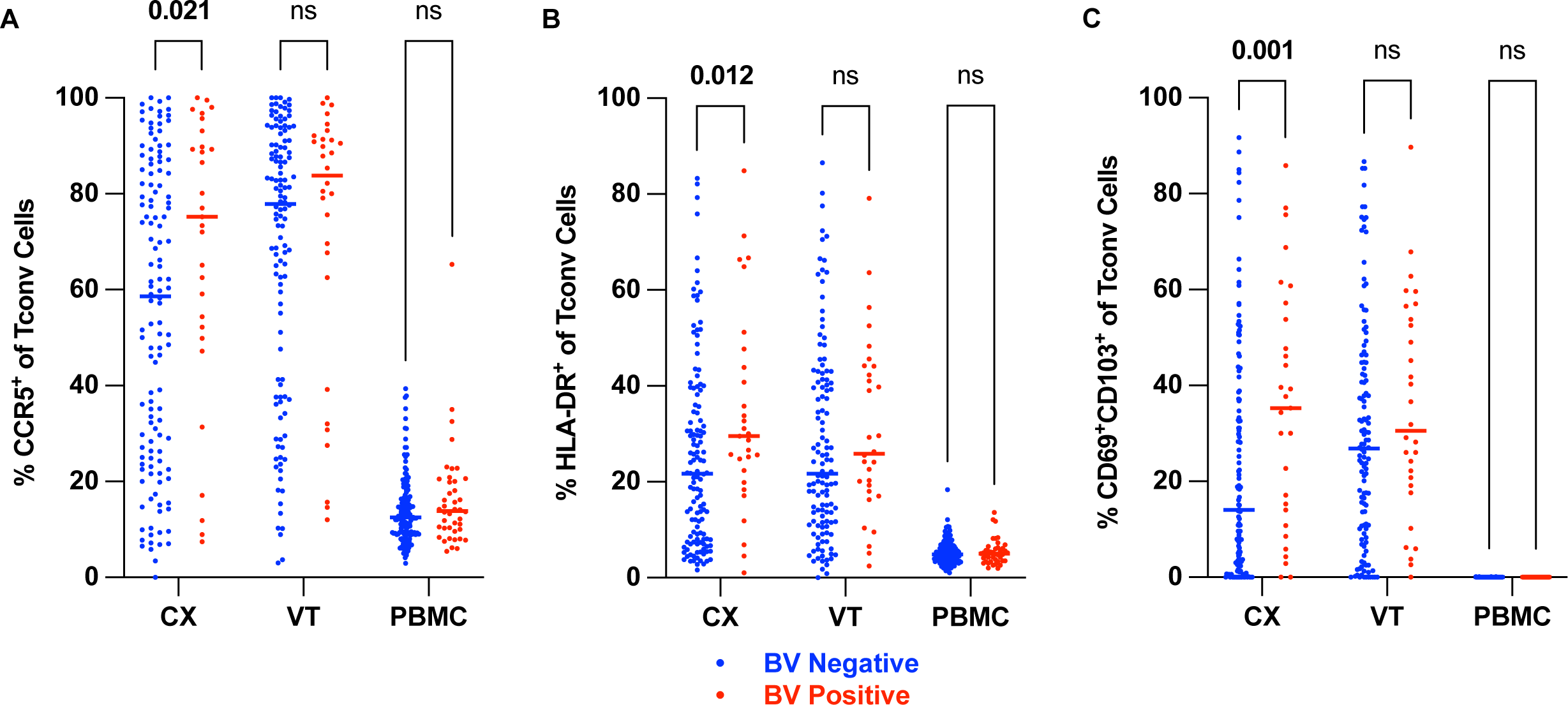
Conventional CD4^+^ T cells displayed increased markers of activation and tissue residency in the cervix of individuals with BV. Flow cytometry was used to examine Tconv cell phenotypes within different tissue sites as indicated. (**A**) The frequency of CCR5^+^, (**B**) HLA-DR^+^, and (**C**) CD69^+^CD103^+^ among the total Tconv T cell population in CX, VT, and PBMC samples in BV- and BV+ individuals. Adjusted rank regression analysis was performed to compare frequencies in each tissue between BV- and BV+ individuals. PBMC comparisons were *a priori* adjusted for hormonal contraceptive use, and CX and VT comparisons were *a priori* adjusted for hormonal contraceptive use, HSV-2 serology, HIV exposure, and semen exposure to reduce the effects of potential confounding variables on the analysis of BV-driven T cell alterations. Adjusted p-value displayed in bold when p ≤ 0.05, non-bold when 0.05 **<** p_adj_ ≤ 0.10, and “ns” for not significant when adjusted p **>** 0.10. Each dot represents a measurement from an individual sample. Each horizontal bar indicates the median for its respective group. For each comparison, the Ns, medians, p-values (adjusted and unadjusted), and estimated differences (adjusted and unadjusted) are provided in Supplemental Table 1.

We hypothesized that T_RM_ would play a critical role in immune modulations associated with BV, so we evaluated CD103^+^CD69^+^ co-expression on CD4^+^ Tconv T cells as canonical T_RM_ markers (24, 46). We found that the T_RM_ phenotype was more frequently observed on CX Tconv cells in BV+ individuals (median frequency 14% BV- vs 35% BV+, p_adj_ = 0.001; **Supplemental Table 1**) and found no significant differences when comparing CD103^+^CD69^+^ Tconv cells isolated from VT or PBMC samples provided by BV- vs BV+ individuals (**Figure 2C**). These results support that BV is associated with an increased proportion of tissue-resident Tconv in the CX but not in the vagina. Not unexpectedly, CD4^+^ T_RM_ were not present in the circulation.

### The total density of T cells and HIV target cells in the cervix and vagina was not altered by BV

To assess whether BV is associated with an increased number or density of CCR5^+^ HIV target T cells in the CVT mucosa, we analyzed VT and CX tissue sections by immunofluorescent staining and microscopy. H&E staining was used to evaluate tissue quality and integrity and visualize the epithelium and lamina propria compartments (representative image **Figure 3A**). Immunofluorescent staining and cellular imaging were used to visualize CD3^+^, CD3^+^CD4^+^, and CD3^+^CD4^+^CCR5^+^ cells in each tissue section (representative image **Figure 3B**). In contrast to the increased frequency of CCR5^+^ Tconv that we observed as a fraction of total Tconv cells in the CX by flow cytometry (**Figure 2A**), we found no significant differences in the overall density of HIV target T cells in the CX (**Figure 3C**, median density 228.3 cells/mm^2^ BV- vs 219.6 cells/mm^2^ BV+, p = 0.764; **Supplemental Table 2**) or VT (**Figure 3D**, median density 227.0 cells/mm^2^ BV- vs 336.2 cells/mm^2^ BV+, p = 0.787; **Supplemental Table 2**). We also observed no difference when analyzing epithelium and lamina propria tissue compartments individually (**Supplemental Figure 2**, **Supplemental Table 2**), suggesting that BV may not significantly alter the overall abundance of HIV target cells within the deeper CX or VT mucosal tissue layers.

**Fig 3.**
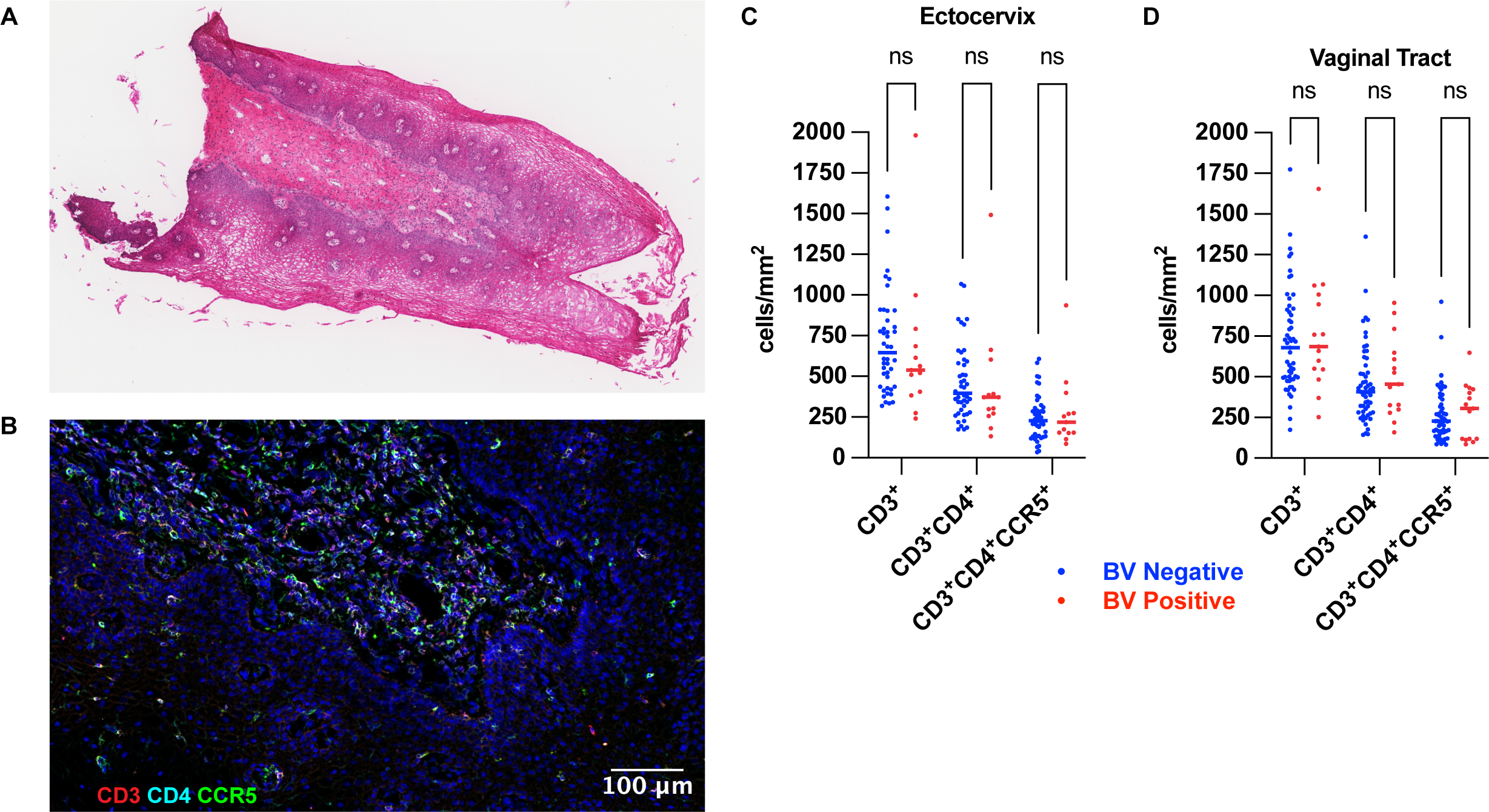
The total density of T cells and HIV target cells in the cervix and vagina was not altered by BV. **(A)** Representative H&E-stained vaginal tract tissue section imaged with brightfield microscopy. **(B)** Representative immunofluorescent stained tissue section from the same vaginal tract (VT) biopsy as Figure 3A. DAPI stain is shown in blue, the CD3 stain is shown in red, the CD4 stain is shown in cyan, and the CCR5 stain is shown in green. All fluorescent signals overlayed. Comparison of the cell density of CD3^+^ T cells, CD3^+^CD4^+^ T cells, or CD3^+^CD4^+^CD5^+^ HIV target cells between BV- and BV+ samples in the ectocervix (**C**) and vaginal tract (**D**). Wilcoxon rank sum test was performed for each comparison shown. Comparisons with p > 0.05 labeled “ns” for not significant. Each dot represents a measurement from an individual sample. Each horizontal bar indicates the median for its respective group. For each comparison, the Ns, medians, p-values (adjusted and unadjusted), and estimated differences (adjusted and unadjusted) are provided in Supplemental Table 2.

### Conventional CD4^+^ T cells in the cervix showed signs of dysfunction in individuals with BV

To more broadly characterize how BV impacts mucosal or systemic T cells that may elucidate alternative immunological mechanisms for adverse outcomes associated with BV, we evaluated Tconv cells for additional markers that could indicate altered functional capacity. We found CD39 expression significantly increased on Tconv cells in CX samples from BV+ individuals (median frequency 11% BV- vs 23% BV+, p_adj_ = 0.005; **Supplemental Table 1**) and in VT samples from BV+ individuals (median frequency 15% BV- vs 23% BV+, p_adj_ = 0.039; **Supplemental Table 1**) with minimal differences observed in the PBMC samples (**Figure 4A**). CD39 is a marker of metabolic stress (53) that may contribute to Tconv dysfunction in the CX and VT of BV+ individuals. CD101 expression was also significantly increased on Tconv cells in CX samples from BV+ individuals (median frequency 19% BV- vs 38% BV+, p_adj_ = 0.002; **Supplemental Table 1**), with minimal differences observed in the VT and PBMC samples (**Figure 4B**). CD101 is another marker associated with tissue residency (23), though it has also been shown to contribute to an inhibitory phenotype (51) that may impede the proliferation of Tconv cells in BV+ individuals. Finally, we found a significant decrease in the frequency of TCF-1^+^ Tconv in the CX of BV+ compared to BV- individuals (median frequency 44% BV- vs 24% BV+, p_adj_ = 0.008; **Supplemental Table 1**), with no significant differences in the VT or PBMC (**Figure 4C**). TCF-1 is a stabilizing transcription factor implicated in the regulation of progenitor potential and promotion of T cell fate specification (54, 59-64). These results suggest that Tconv cells in the CX of BV+ individuals may have a reduced capacity to undergo differentiation or serve as proliferative progenitor cells, which, taken together, could suggest that these cells have reduced capacity to serve as potent effector T cells.

**Figure 4.**
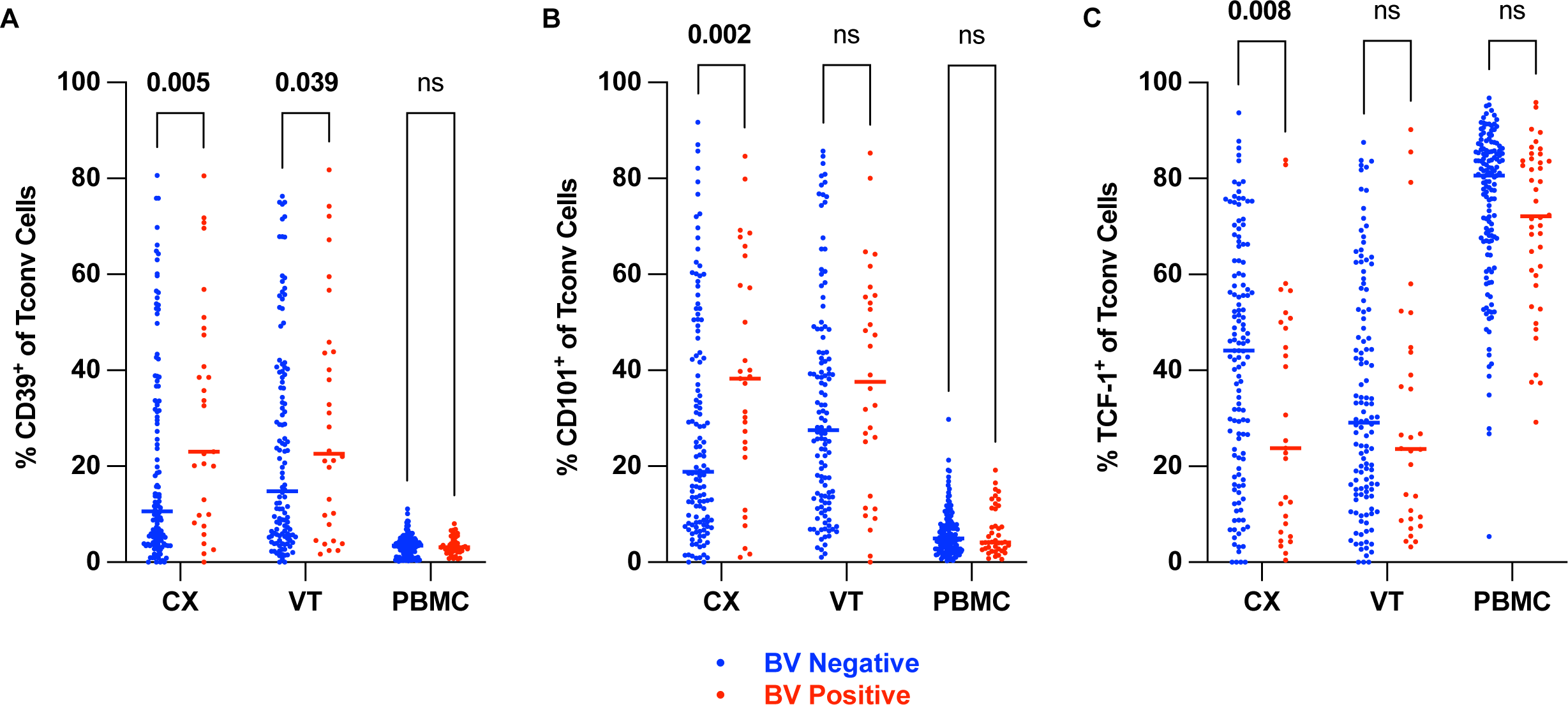
Conventional CD4^+^ T cells in the cervix showed signs of dysfunction in individuals with BV. Flow cytometry was used to examine Tconv cell phenotypes within different tissue sites as indicated. (**A**) The frequency of CD39^+^, (**B**) CD101 ^+^, and (**C**) T Cell Factor-1^+^ (TCF-1) among the total Tconv T cell population in CX, VT, and PBMC samples in BV- and BV+ individuals. Adjusted rank regression analysis was performed to compare frequencies in each tissue between BV- and BV+ individuals. PBMC comparisons were *a priori* adjusted for hormonal contraceptive use, and CX and VT comparisons were *a priori* adjusted for hormonal contraceptive use, HSV-2 serology, HIV exposure, and semen exposure to reduce the effects of potential confounding variables on the analysis of BV-driven T cell alterations. Adjusted p-value displayed in bold when p ≤ 0.05, non-bold when 0.05 **<** p_adj_ ≤ 0.10, and “ns” for not significant when adjusted p **>** 0.10. Each dot represents a measurement from an individual sample. Each horizontal bar indicates the median for its respective group. For each comparison, the Ns, medians, p-values (adjusted and unadjusted), and estimated differences (adjusted and unadjusted) are provided in Supplemental Table 1.

### Th17 cells exhibited increased markers of activation and tissue residency in cervical samples from individuals with BV

Lineage-specific Th17 Tconv (Th17) cells have been shown to play a role in immune responses to extracellular bacteria (65-67). To investigate the impact of BV on Th17 cell phenotypes, we first analyzed the frequency of Th17 cells (defined as CD161^+^CCR6^+^ Tconv cells (49); **Supplemental Figure 1**) among total Tconv cells and found that the frequency of Th17 cells as a proportion of total Tconv cells was not significantly increased in any of the tissue types for BV+ compared to BV- individuals (**Figure 5A**). However, the frequency of Th17 cells expressing the proinflammatory activation marker HLA-DR was significantly increased in CX during BV (median 40% BV- vs 55% BV+, p_adj_ = 0.024, **Supplemental Table 1**) with a trend towards an increase in the VT (median 38% BV- vs 54% BV+, p_adj_ = 0.067, **Supplemental Table 1**) and with minimal differences in PBMC (**Figure 5B**).

**Figure 5.**
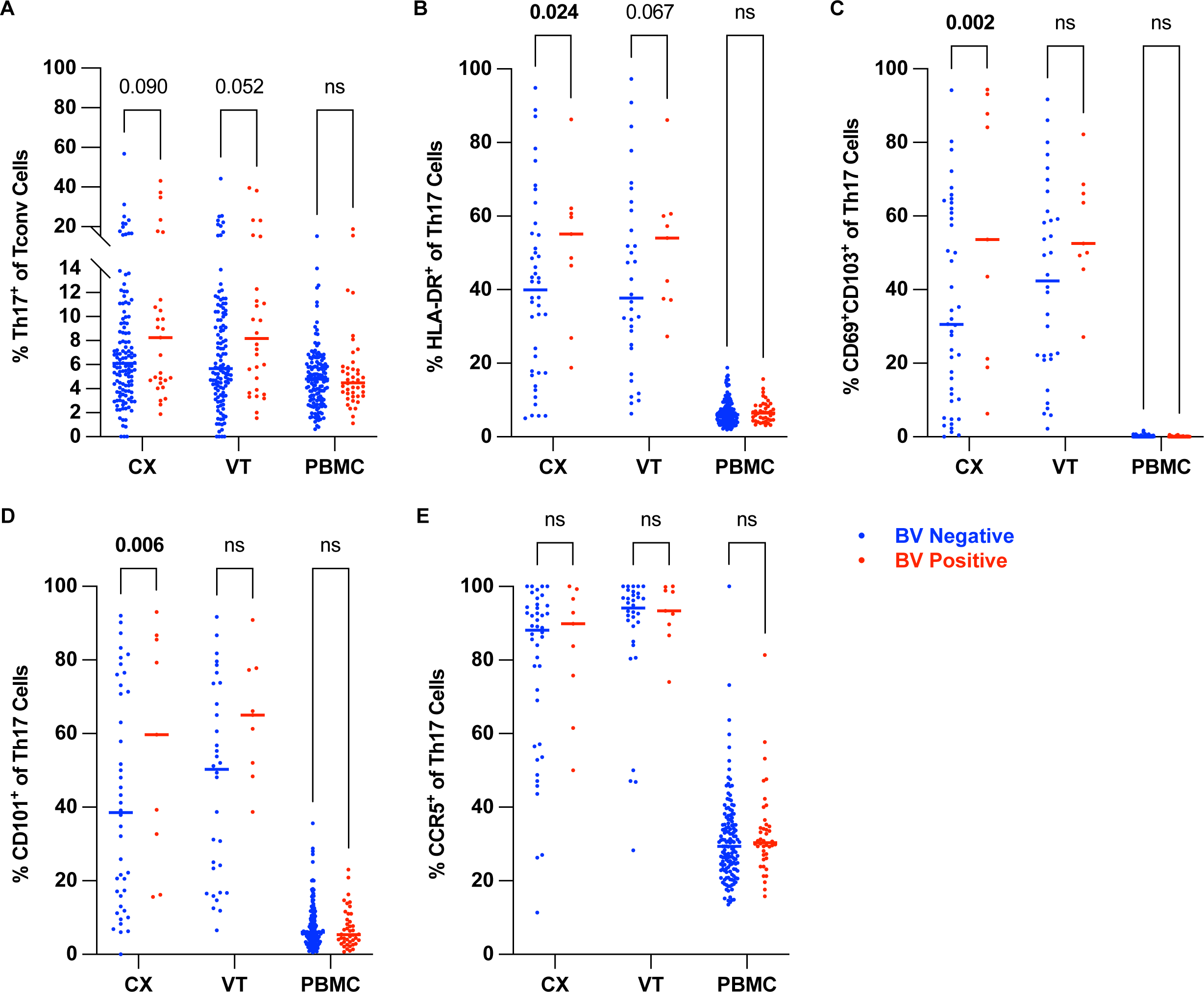
Th17 cells exhibited increased markers of activation and tissue residency in cervical samples from individuals with BV. Flow cytometry was used to examine Th17 phenotypes within different tissue sites as indicated. **(A)** The frequency of Th17 T cells, defined as CD161^+^CCR6^+^ Tconv, among the total Tconv T cell population in CX, VT, and PBMC samples in BV- and BV+ individuals. (**B**) HLA-DR ^+^, (**C**) CD69^+^CD103^+^, (**D**) CD101 ^+^, and (**E**) CCR5^+^ frequencies among the total Th17 population in CX, VT, and PBMC samples in BV- and BV+ individuals. Adjusted rank regression analysis was performed to compare frequencies in each tissue between BV- and BV+ individuals. PBMC comparisons were *a priori* adjusted for hormonal contraceptive use, and CX and VT comparisons were *a priori* adjusted for hormonal contraceptive use, HSV-2 serology, HIV exposure, and semen exposure to reduce the effects of potential confounding variables on the analysis of BV-driven T cell alterations. Adjusted p-value displayed in bold when p ≤ 0.05, non- bold when 0.05 **<** p_adj_ ≤ 0.10, and “ns” for not significant when adjusted p **>** 0.10. Each dot represents a measurement from an individual sample. Each horizontal bar indicates the median for its respective group. For each comparison, the Ns, medians, p-values (adjusted and unadjusted), and estimated differences (adjusted and unadjusted) are provided in Supplemental Table 1.

We also measured the frequency of markers of tissue-residency (CD69^+^CD103^+^) on Th17 cells and found them significantly increased in the CX of BV+ compared to BV- individuals (median 31% BV- vs 54% BV+, p_adj_ = 0.002, **Supplemental Table 1**) (**Figure 5C**). In addition, the frequency of Th17 cells expressing CD101, consistent with a T_RM_ phenotype, was significantly increased in the CX of BV+ compared to BV- individuals (median 39% BV- vs 60% BV+, p_adj_ = 0.006, **Supplemental Table 1**) (**Figure 5D**).

Finally, we found that the frequency of Th17 cells expressing the activation marker and HIV co-receptor CCR5 was not significantly different between BV- vs BV+ individuals, although notably, the median frequency of Th17 cells expressing CCR5 in BV+ individuals was 90% in CX and 93% in VT samples (**Supplemental Table 1**) (**Figure 5E**). Therefore, CVT Th17 cells predominantly express the HIV coreceptor CCR5, coinciding with previous reports that Th17 cells are viable targets for HIV infection (68). These results demonstrate that BV is associated with CX Th17 activation and that the increased tissue-resident or CD101-expressing CX Th17 cells may contribute to a mechanism for increased HIV susceptibility in individuals with BV.

### Cervical CD8^+^ T cells displayed a dysfunctional phenotype in individuals with BV

CD8^+^ T cells are thought to play an important role in pathogen clearance in the CVT, including against BV-associated organisms. To comprehensively evaluate how BV impacts CD8^+^ T cell phenotypes in the CVT mucosa and circulation, we analyzed CX, VT, and PBMC samples for the expression of effector markers (**Supplemental Figure 1** and **Supplemental Table 1**). We found that the frequency of CD39^+^CD8^+^ T cells was significantly increased in CX samples from BV+ individuals (median frequency 7% BV- vs 10% BV+, p_adj_ = 0.023; **Supplemental Table 1**), and moderately but insignificantly increased in the VT of BV+ individuals (median frequency 6% BV- vs 10% BV+, p_adj_ = 0.068; **Supplemental Table 1**, **Figure 6A**). We also observed a moderate though insignificant decrease in Granzyme B expression, a marker of cytotoxicity, on CD8^+^ T cells in the CX of BV+ individuals (median frequency 25% BV- vs 22% BV+, p_adj_ = 0.094; **Supplemental Table 1**) with no significant differences in the VT and PBMC samples from those with vs without BV (**Figure 6B**).

**Figure 6.**
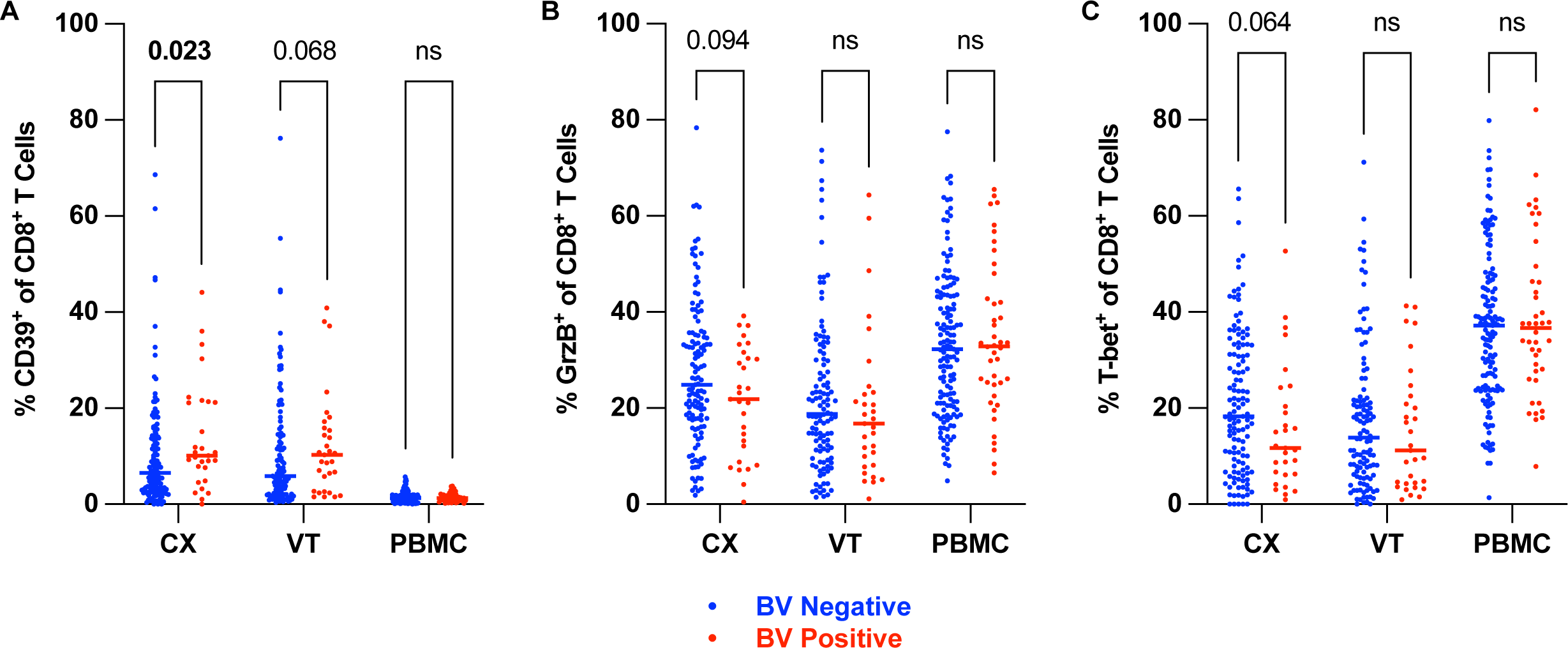
Cervical CD8^+^ T cells displayed a dysfunctional phenotype in individuals with BV. Flow cytometry was used to examine CD8^+^ T cell phenotypes within different tissue sites as indicated. (**A**) The frequency of CD39^+^, (**B**) Granzyme B^+^, and (**C**) T-bet^+^ among the total CD8^+^ T cell population in CX, VT, and PBMC samples in BV- and BV+ individuals. Adjusted rank regression analysis was performed to compare frequencies in each tissue between BV- and BV+ individuals. PBMC comparisons were *a priori* adjusted for hormonal contraceptive use, and CX and VT comparisons were *a priori* adjusted for hormonal contraceptive use, HSV-2 serology, HIV exposure, and semen exposure to reduce the effects of potential confounding variables on the analysis of BV-driven T cell alterations. Adjusted p-value displayed in bold when p ≤ 0.05, non-bold when 0.05 **<** p_adj_ ≤ 0.10, and “ns” for not significant when adjusted p **>** 0.10. Each dot represents a measurement from an individual sample. Each horizontal bar indicates the median for its respective group. For each comparison, the Ns, medians, p-values (adjusted and unadjusted), and estimated differences (adjusted and unadjusted) are provided in Supplemental Table 1.

Additionally, we observed a moderate but insignificant decrease in the frequency of T-bet^+^ CD8^+^ T cells in the CX of BV+ individuals (median frequency 18% BV- vs 12% BV+, p_adj_ = 0.064; **Supplemental Table 1**) with no significant differences in the VT and PBMC samples from those with vs without BV (**Figure 6C**). Altogether, these findings suggest a trend towards decreased CX CD8^+^ T cell activation in addition to an increased dysfunctional CD39^+^ CD8^+^ T cell phenotype observed in the cervix of BV+ individuals.

### BV was associated with reduced chemokine concentrations and increased inflammatory cytokine concentrations in CVT fluid

To investigate soluble immune factors in individuals with BV versus those without BV, we analyzed 71 cytokines and chemokines in serum and CVT fluid collected via menstrual cup. In CVT fluid samples, 61 cytokines and chemokines met the criteria for continuous analysis (**Figure 7A**), while 10 cytokines and chemokines were analyzed as a dichotomous outcome (detected/not detected; **Supplemental Figure 3A**). In the serum, 52 different cytokines and chemokines met the criteria for continuous analysis (**Figure 7B**), 17 cytokines and chemokines were analyzed using the dichotomous outcome (**Supplemental Figure 3B**), and 2 cytokines were detectable in all individuals but had greater than 20% of samples out of range high so were excluded from the analysis (**Supplemental Table 3**). We observed significantly higher concentrations of twenty-three soluble immune factors including IL-1α in the menstrual cup fluid collected from participants with versus without BV (**Figure 7A**). Additionally, we observed significantly lower concentrations of twelve soluble immune factors, including MIG/CXCL9 and IP-10 (CXCL10) (**Figure 7A**) in participants with BV versus those without BV. In our analysis of serum cytokines, fewer significant observations were observed than in the CVT fluid. In total, we observed a significant increase of one and a significant reduction of five quantifiable soluble immune factors in participants with BV (**Figure 7B**). In the CVT fluid samples, of the ten soluble immune factors analyzed for their dichotomous outcome, three were detected significantly more frequently, and one cytokine was detected significantly less frequently in BV+ individuals (**Supplemental Figure 3A**). The dichotomous outcome analysis revealed an additional two other cytokines observed significantly less frequently in the serum of BV+ individuals (**Supplemental Figure 3B**). Overall, differentially expressed and detected soluble immune factors were primarily observed in CVT fluid samples, with fewer differences observed in serum samples, highlighting that BV has the greatest effects on the local CVT, as opposed to the circulating immune environment.

**Figure 7.**
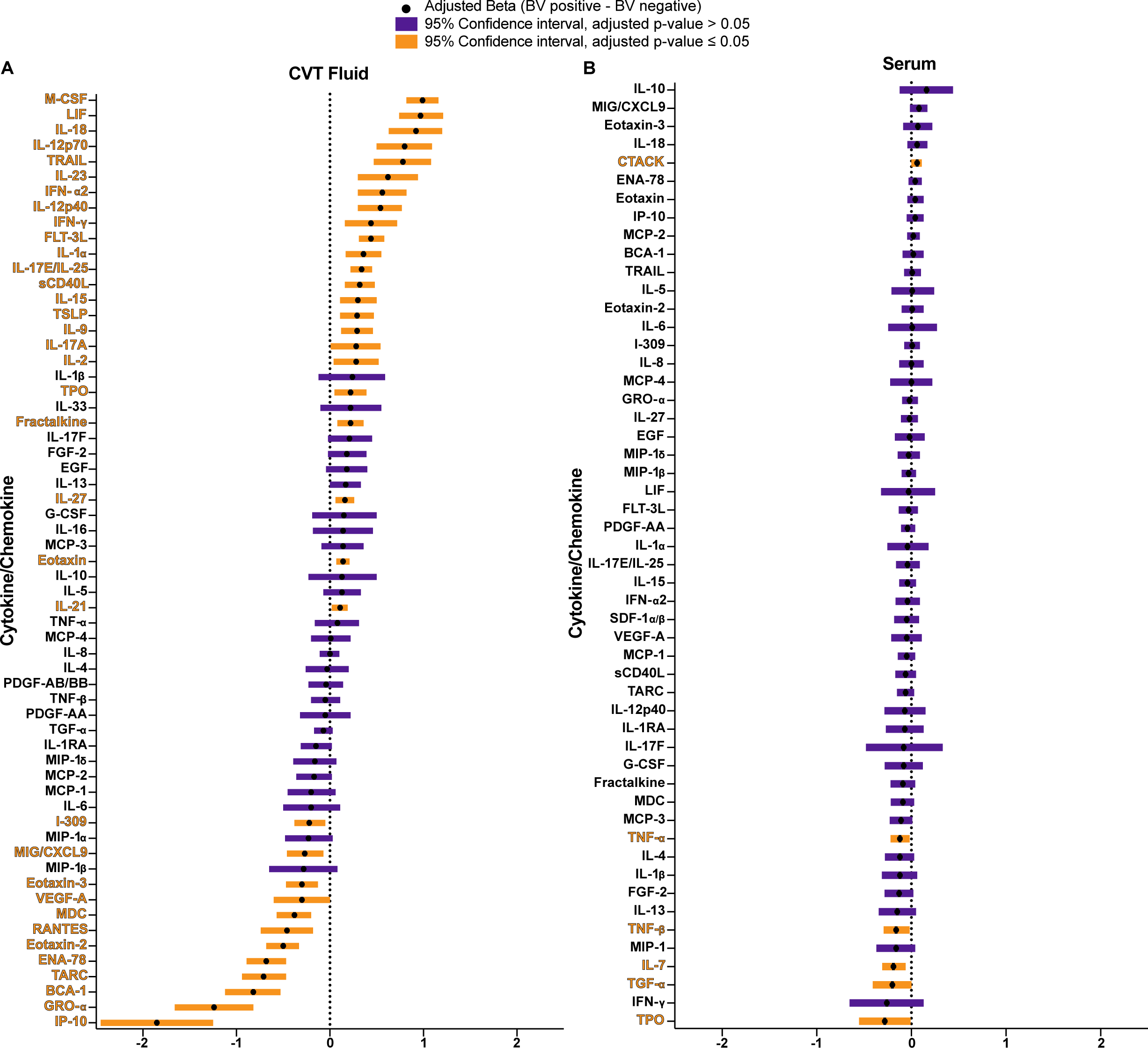
BV was associated with reduced chemokine concentrations and increased inflammatory cytokine concentrations in CVT fluid. Luminex was used to quantify cytokines and chemokines from CVT fluid or serum samples. Adjusted estimated difference (BV positive – BV negative) for CVT fluid cytokines (**A**) and serum (**B**) that met the criteria for quantification of cytokine concentrations (greater than or equal to 80% of samples were detectable). The adjusted 95% confidence interval is shown for all comparisons. Significant results when adjusted p ≤ 0.05 comparing BV- vs BV+ are colored in orange and non-significant differences (p > 0.05) comparing BV- vs BV+ are purple. Vertical dashed line at x=0 for reference. Serum comparisons were *a priori* adjusted for hormonal contraceptive use, and CVT fluid comparisons were *a priori* adjusted for hormonal contraceptive use, HSV-2 serology, HIV exposure, and semen exposure to reduce the effects of potential confounding variables on the analysis of BV-driven T cell alterations. Statistical analyses can be found in Supplemental Table 3.

## Discussion

In context of the global HIV pandemic, T cells serve as a double-edged sword: mediating anti-viral functions required to fight the virus while also acting as potential targets for HIV infection and replication (69, 70). The findings presented here add further detail that may underlie the effect of BV on HIV susceptibility by highlighting a more complex scenario that involves deleterious effects on T cell antiviral functions in addition to modulations in subsets that may act as potential HIV target cells.

Our flow cytometry analysis revealed an increase in the frequency of activation markers expressed on cervical Tconv, including a higher frequency of CCR5^+^ among total Tconv cells (**Figure 2A**), which supports prior studies that have identified increased levels of total CD3^+^CD4^+^CCR5^+^ T cells in the CVT lumen as a potential source of HIV target cells (7, 39, 40). However, our analysis of CD3^+^CD4^+^CCR5^+^ HIV target cell tissue density by immunofluorescent staining and cellular imaging of tissue sections indicated a relatively equivalent abundance of CD3^+^CD4^+^CCR5^+^ HIV target cells among BV- and BV+ participants in CX and VT tissue evaluated overall (**Figure 3, C** and **D**) and when the epithelium and lamina propria compartments were analyzed separately (**Supplemental Figure 2**). This supports the idea that BV does not have a profound effect on the overall abundance of HIV target cells in the cervicovaginal mucosa. Our conclusions are limited by the fact that the results are based in distinct technologies (flow cytometry versus microscopy); and that flow cytometry involves the evaluation of more samples and therefore has increased power to detect differences compared to microscopy. Nevertheless, our results underscore that the abundance of CD3^+^CD4^+^CCR5^+^ HIV target cells within the deeper CX and VT tissue layers in BV+ individuals is unlikely to solely account for increased HIV susceptibility among those with BV, and this motivates the possibility that other factors contribute to this adverse outcome.

Our application of high-parameter, high-throughput flow cytometry on cells isolated from VT and CX tissue biopsies and PBMC samples documents that BV elicits complex changes to CVT T cells that may contribute to the inhibition of antiviral T cell function. We observed an increased frequency of CD4^+^ Tconv cells expressing CD39 in the CX and VT of BV+ individuals (**Figure 4, A and B**). The potential dysfunction of this T cell phenotype is suggested by prior studies showing CD39^+^ T cells have a decreased response to vaccines and an increased likelihood of undergoing apoptosis (53), raising the possibility that in the context of BV, cervical and vaginal CD4^+^ Tconv may exhibit increased intrinsic immunoregulation leading to reduced effector potential. This could lead to increased HIV susceptibility through an inability to mount an effective antiviral response upon exposure to the virus.

We also found that CD101^+^ Tconv cells, and specifically CD101^+^ Th17 cells, are more frequently observed in the CX of BV+ individuals (**Figure 4B and 5D**). Less is known about the function of CD101^+^ T cells in comparison to the other phenotypes examined in this study, but CD101^+^ T cells have been associated with decreased expansion after adoptive transfer in mice (51), suggesting CD101^+^ Tconv cells are inhibited and less likely to undergo proliferation. Additionally, CD101 is associated with exhausted T cell phenotypes that result from chronic activation and can promote immune dysfunction (71-73). Thus, the increased abundance of CD101^+^ Tconv and Th17 cells in the CX of BV+ individuals may also contribute to a dysfunctional Tconv immune response upon secondary exposure to HIV in the CX. Given the often recurrent and persistent nature of BV (44), chronic immune activation from repeated antigen exposure caused by BV-associated microbiota may lead to the progression of these observed dysfunctional phenotypes that could promote HIV susceptibility by limiting antiviral T cell function, expansion, and differentiation.

We also observed a significant reduction in the proportion of TCF-1^+^ CD4^+^ Tconv in the CX of BV+ individuals (**Figure 4C**). TCF-1 is necessary for follicular T cell differentiation (59-61) and has been shown to play a role in Th17 responses by inhibiting IL-17 production (64). Fewer progenitor CD4^+^ T cells implies that more T cells are likely to be terminally differentiated in the CX, which may further limit lineage-specific differentiation upon secondary exposure to an infectious agent.

In addition to CD4^+^ T cells expressing CCR5, it has been shown that CD69^+^CD4^+^ T cells are associated with increased susceptibility to HIV infection in the CVT mucosal tissue (74). This suggests that the increased frequency of T_RM_ in the CX of BV+ individuals (**Figure 2C**) may mediate increased HIV acquisition in the context of BV. In particular, we found the CX had an increased frequency of Th17 T_RM_ (**Figure 5C**), which highly expresses CCR5 (**Figure 5E**). The possibility that these cells are a target for HIV acquisition in BV+ individuals is supported by a prior report that Th17 cells serve as primary target cells during vaginal SIV infection (75). In addition to the role CD4^+^ T_RM_ in CVT mucosa play in HIV susceptibility, they also act as reservoirs for HIV viral replication which can occur during early HIV infection (76). The increased frequency of Tconv T_RM_, and in particular Th17 T_RM_, may not only contribute to susceptibility to primary HIV infection, but may also promote viral replication in the CX of BV+ individuals.

Activated CD8^+^ T cells are thought to be important in controlling HIV replication after initial infection (77, 78). The increased frequency of mucosal CD39^+^ CD8^+^ T cells observed in individuals with BV may also augment CVT immune dysfunction. Trends toward decreased CX Granzyme B^+^ or T-bet^+^ CD8^+^ T cell activation (**Figure 6B and C**) may relate to the increased frequency of dysfunctional CD4^+^ Tconv cells observed (**Figures 4 and 5**), since functional CD4^+^ Tconv cells are known to enhance CD8^+^ T cell effector function (79), and in the context of BV, CD4+ Tconv cells appear to be diminished in function. The interactions, or possibly lack thereof, between CD8^+^ T cells and dysfunctional Tconv cells in the context of BV, may also contribute to a dysfunctional CD8^+^ T cell response during BV and increased HIV susceptibility in BV+ individuals by inhibiting an effective CD8^+^ T cell response after exposure to HIV.

Our analysis of soluble immune factor detection in BV+ vs BV- individuals further demonstrates the diverse immune modulations that BV can promote in the CVT. We observed significant increases in many pro-inflammatory cytokines in BV+ CVT fluid samples, including IL-1α, as previously reported by other studies (80-82). We also observed significantly increased IL-17A, IFN-γ, IL-21, IL-23, and IL-12p70 in the CVT fluid of BV+ individuals (**Figure 7A**), all of which can each enhance Th17-mediated inflammation (49, 66, 83-85). The activation of Th17 cells may then lead to the formation of Th17 T_RM_ which were observed more frequently in the CX of BV+ individuals by flow cytometry (**Figure 5C**). Additionally, increased M-CSF in the CVT fluid of BV+ individuals (**Figure 7A**), produced by macrophages, has been shown to promote Th17 differentiation of CD4^+^ memory T cells (86), which may also contribute to the expanded Th17 T_RM_ pool in the CX of BV+ individuals. On the other hand, LIF and IL-17E/IL-25, observed to be increased in the CVT fluid of BV+ individuals (**Figure 7A**), inhibit inflammatory responses, including the inhibition of Th17-mediated immune responses (87, 88). LIF and IL-17E/IL-25 may be produced in response to the activation of Th17 cells to limit overall Th17 cell differentiation and proliferation. The reduction of TCF-1^+^ progenitor Tconv cells concurrent with the production of LIF and IL-17E/IL-25 in the CVT could limit the differentiation of progenitor T cells to Th17 cells, which could explain why we do not observe a significant overall expansion in frequency of Th17 cells in BV+ individuals (**Figure 5A**).

While we observe an increased concentration of many proinflammatory cytokines in BV+ CVT fluid samples, we also observe the reduction of many T cell recruiting chemokines. Our results support previous studies that have also observed reductions of IP-10 (CXCL10) and MIG/CXCL9 in the CVT of BV+ individuals (82, 89-91). We hypothesized that RANTES (CCL5) would be increased in BV+ individuals (37) to promote the recruitment of CCR5^+^ HIV target cells to the CVT. However, our observed reduction of RANTES in the CVT fluid of BV+ individuals has also been reported in a prior study that evaluated CVT fluid collected by menstrual cup (90). While RANTES does act as a primary ligand for CCR5, which can promote the recruitment of CCR5^+^ cells, it also reduces cell-surface expression of CCR5 by binding and internalizing the CCR5 receptor (92). Increased CVT RANTES has been associated with increased HIV resistance (93), possibly by competing with HIV for CCR5 binding (94). Reduced RANTES in BV+ individuals may limit the number of CCR5^+^ HIV target cells that are recruited to the CVT. Simultaneously, this could promote stabilization of CCR5 expression on existing T cells in the CVT by reducing ligand availability and binding to internalize residually expressed CCR5, a protein preferentially observed on CVT Tconv cells vs circulating Tconv cells (**Figure 2A**). The relationship between CCR5 expression and CVT RANTES production in BV+ individuals is complex and how these factors impact HIV susceptibility over time will require more detailed longitudinal observational studies. The reduction of IP-10, MIG/CXCL9, RANTES, and several other proinflammatory chemokines in the CVT fluid of BV+ individuals supports the notion of impaired T cell recruitment to the CVT, which adds further evidence to dysregulated CVT immune responses in BV+ individuals. This may also contribute to adverse health outcomes, including increased HIV susceptibility by inhibiting antiviral T cell recruitment to the CVT in BV+ individuals.

In addition to our *a priori* hypothesis of an increase in RANTES in the CVT fluid of BV+ individuals, we also hypothesized that IL-6, IL-8, and IL-1β would each be increased in BV+ individuals based on a previous meta-analysis (37). We observed no significant difference in IL-6, IL-8, or IL-1β in CVT samples from BV+ vs BV- individuals. However, CVT IL-1β was trending towards an increase in BV+ (adjusted log mean difference BV+ - BV- = 0.24; p_adj_ = 0.188; **Figure 7A, Supplemental Table 3**). While IL-1β is frequently described as increased in CVT fluid from BV+ individuals, this is not a universal observation (95, 96), nor is IL-6 or IL-8 uniformly increased across cohorts of BV+ individuals (97, 98). The factors that mediate altered expression of soluble immune mediators during BV have not been elucidated, highlighting the complexity and variability of BV-driven CVT immune responses.

A majority of the significant phenotypic alterations observed in this study occurred when comparing CX samples from individuals with vs without BV. This indicates that while BV is a dysbiosis of vaginal flora, the immune cells in the CX may be more greatly impacted by exposure to BV-associated organisms than the VT. One reason for this may be that the unique CX microbiota (99) may be more sensitive to changes in flora than the VT, or that the immune cells of the VT may be more resistant to BV-driven phenotypic alterations than those in the CX. The primary cellular targets for intravaginal infection of SIV in rhesus macaques are in the CX lamina propria (100), suggesting that the CX may be the primary site of sexual transmission of HIV in humans. Therefore, the more distinct immune alterations observed in the CX of BV+ individuals may be of particular concern for the increased risk of HIV transmission and replication in BV+ individuals. Comparing immune modulations in the CX vs VT among those with or without BV warrants further exploration.

Our study has several limitations: first, we focused on characterizing diverse T cell markers and a large panel of soluble immune mediators in the context of BV infection, leaving the relationship of BV to other components of the immune response (monocytes/macrophages, granulocytes, and other immune cell types) unevaluated. One justification for limiting our evaluation to characteristics of CD3^+^ T cells is that these constitute the predominant lymphocyte population in the CVT mucosa (**Figure 1A**). A second limitation was that we collected 3-mm tissue biopsies that often captured low amounts of rare T cell subsets such as Tregs. We were underpowered to evaluate CX Treg phenotypes (N BV+ = 2) or VT Treg phenotypes (N BV+ = 4) (data not shown). This also reduced our ability to effectively evaluate the role of other rare subsets in immune responses to BV. Third, an alternative hypothesis for adverse health outcomes associated with BV is that BV drives epithelial damage which, in the context of HIV susceptibility, reduces the effectiveness of the epithelial barrier in providing a physical barrier to protect against HIV transmission. We attempted to quantify tissue damage (data not shown) but were limited by sample quality for blinded pathology scoring. A fourth limitation to our study was that tissue biopsies were variable in size, and so we cannot compare cell numbers in tissue by flow cytometry. Future studies will include weighing tissue biopsies prior to cryopreservation so that we can normalize the number of cells analyzed by flow cytometry per tissue biopsy by weight. A final limitation of our analysis is that it was performed as an exploratory, hypothesis-generating analysis, and as such, used nominal P values adjusted for confounders but not discounted for multiple comparisons (138 immune cell subsets comparisons, and 71 soluble mediator comparisons). Thus, our findings should be confirmed through studies employing hypothesis-driven testing.

In summary, our study performed the most comprehensive evaluation to date of CVT tissue T cell subsets associated with BV. This data shows BV-driven immune alterations have the most profound impact on CX T cells and changes at this site may be most responsible for increased HIV susceptibility in BV+ individuals. While we observe increased CX Tconv activation including increased frequency of CCR5 detection on CX Tconv cells by flow cytometry, our analysis of CX and VT tissue sections by immunofluorescent microscopy shows that BV does not have a profound effect on overall CCR5^+^ HIV target cell density. Our high-parameter, high throughput flow cytometry analysis revealed that BV drives diverse phenotypic alterations that extend beyond CD4^+^ Tconv activation in the CVT. Specifically, we identify an increased frequency of dysfunctional T cell subsets, including CD39^+^ Tconv and CD39^+^ CD8^+^ T cells, and a decreased frequency of progenitor TCF-1^+^ Tconv in the CX of BV+ individuals that could alter host response and contribute to increased HIV susceptibility by limiting the antiviral capabilities of CX T cells. Furthermore, we found the enrichment of Tconv T_RM_, and specifically the Th17 T_RM_ subset, in the CX of BV+ individuals that may also serve as targets for HIV infection and replication. Confirmation of these findings and elaboration on molecular mechanisms may identify novel targets for immune interventions to reduce the risk of adverse health outcomes associated with BV, including increased risk of HIV infection.

## Methods

### Participants, samples, and data collection

Samples and data for this analysis came from the Kinga Study (Clinicaltrials.gov ID# NCT03701802) which enrolled a total of 406 heterosexual Kenyan couples from Oct. 2018 through Dec. 2019 to evaluate how exposure to sexually transmitted infectious agents alters genital mucosal immune responses. Among these couples, 110 were HIV serodifferent (defined as a person living with HIV (PLWH) and their heterosexual HIV-exposed partner), with the remaining 298 couples involving partners who were both HIV-seronegative at enrollment. HIV serodifferent couples were excluded if, prior to enrollment, the PLWH had initiated antiretroviral therapy (ART) with resulting suppressed HIV viral load, or the HIV-exposed partner had initiated tenofovir-based pre-exposure prophylaxis (PrEP). ART and PrEP were provided to enrolled PLWH or HIV-exposed partners, respectively, upon enrollment.

Seven sample types were requested from all Kinga Study participants at enrollment and 6-month follow-up visits for evaluation of immune responses: cervicovaginal tract fluid via Softcup^®^ (menstrual cup); two 3-mm vaginal tissue biopsies with one cryopreserved for immunofluorescent imaging, and one cryopreserved for flow cytometry; two 3-mm ectocervical tissue biopsies processed in parallel to the vaginal biopsies; fractionated peripheral blood mononuclear cells; and serum were all collected (Note: genital samples and CVT fluid collection were deferred if participants were actively menstruating and participants were encouraged to return to clinic when not menstruating for collection of all sample types at the same timepoint). Swabs for BV testing were collected prior to biopsies, and BV was assessed by Nugent score with 0-3 defined as normal flora, 4-6 as intermediate flora, and 7-10 classified as BV (101). Additional demographic, epidemiologic, clinical, and sexual behavior data (including self-reported frequency of vaginal sex, condom use, and hormonal contraceptive use) were collected at all visits.

Laboratory testing at the enrollment and 6-month follow-up visits included HIV serology by Determine HIV 1/2 Rapid diagnostic Test (RDT) (Abbott Laboratories, Inc., Abbott Park, IL, USA) with confirmation of positive results with First Response RDT (Premier Medical Corporation Ltd., Kachigam, India), and further confirmation of RDT results using 4th generation HIV-1-2 Ag/Ab Murex EIA assay (DiaSorin, Inc., Kent, UK). Herpes simplex virus type 2 (HSV-2) serology was performed with HerpeSelect 2 ELISA IgG test (Focus Technologies, Inc., Cypress, CA) using an index value cut-off of ≥3.5 to improve test specificity (102-106).

To identify samples for the current cross-sectional analysis of the effects of BV on immune responses we focused on 245 participants living without HIV who were identified as female at birth. These consisted of all 44 participants who may have been exposed to HIV by their enrolled heterosexual sexual partners, and 201 whose enrolled heterosexual sexual partner was without HIV and whose data could support a variety of analyses, including the analysis of BV mediated immune responses. Immunofluorescent imaging was performed on enrollment samples from a subset who were HSV-2 seronegative and had HIV-uninfected partners. All immunologic testing was performed blinded to participant exposure data.

### Analysis of tissue samples by high-parameter flow cytometry

*Sample collection:* Biopsies were collected using baby Tischler forceps biopsies either at the lateral vaginal wall or the ectocervical os, placed in a cryovial containing fetal bovine serum at 4°C, and transported to the lab. In the lab, dimethyl sulfoxide (DMSO) was added to the cryovial to a final concentration of 10%, cryopreserved overnight at -80°C, and transferred to liquid nitrogen for long-term storage as described previously by Hughes *et al* (45).

Peripheral blood mononuclear cells (PBMCs) were isolated from acid-citrate dextrose (ACD) whole blood by centrifugation and resuspension in Dulbecco’s Phosphate Buffered Saline (DPBS). Cells were centrifuged over Ficoll-Histopaque (Sigma-Aldrich, Cat# 10771) at room temperature. The buffy coat was collected and washed twice in DPBS at 4°C before live cells were counted on a hemacytometer with trypan blue. Cells were pelleted and resuspended in 10% DMSO in fetal calf serum at 4°C at a concentration of either 5-7 x 10^6^ cells/mL, or 10-15 x 10^6^ cells/mL. Cryovials with 1 mL of these PBMC suspensions were placed in controlled rate freezing equipment (Mr. Frosty) and stored overnight at -80° C. Cells were subsequently transferred to liquid nitrogen for long term storage and shipped in liquid nitrogen dry shippers to University of Washington for further analysis.

*Tissue Processing for Flow Cytometry:* Cryopreserved PBMCs, ectocervical biopsies, or vaginal biopsies in liquid nitrogen were quickly thawed and transferred to 10% FBS (PBMCs) or 7.5% FBS (VT/CX biopsies) complete RPMI media (RP10 or RP7.5). PBMCs were spun down and resuspended in RP10. After sitting in warmed RP7.5 for 10 minutes, biopsies were incubated at 37 degrees C for 30 minutes with Collagenase II (Sigma-Aldrich, 700 U/mL) and DNase I (Sigma-Aldrich, 400U/mL) in RP7.5. Biopsies were then passed through a 100-mm cell strainer using a plunger to disrupt the tissue, washed, and resuspended in phosphate-buffered saline (PBS). PBMCs and cells isolated from tissues were stained immediately after processing for flow cytometry. Cryopreserved PBMCs from a healthy control donor (Seattle Area Control Cohort (SAC)) were used as a longitudinal technical control for all flow cytometry acquisitions (data not shown).

*Flow Cytometry:* Immediately following isolation, cells were incubated with UV Blue Live/Dead reagent (**Supplemental Table 4**) in PBS for 30 mins at room temperature. After washing, cells were then stained with biotinylated CXCR3 (**Supplemental Table 4**) in 0.5% FACS buffer for 20 mins at room temperature. Cells were washed and then stained extracellularly with antibodies (**Supplemental Table 4**) diluted in 0.5% FACS buffer and brilliant staining buffer (BD Biosciences, Cat# 563794) for 20 mins at room temp. Cells were fixed with Foxp3 Transcription Factor Fixation/Permeabilization buffer (ThermoFisher, Cat# 00-5521-00) for 30 mins at room temp. Cells were washed and then stained intracellularly with antibodies diluted in 1X Permeabilization Buffer (ThermoFisher, Cat# 00-8333-56) for 30 mins at room temp. Cells were then resuspended in 200 ml 0.5% FACS buffer and stored at 4 degrees until ready for use. Antibodies were titrated and used at optimal dilution. Staining was performed in 5-ml polystyrene tubes (Falcon, Cat# 352054). Analysis was performed using Flowjo software and we required at least 25 cells for analysis of phenotypes associated with the parent cell type and further downstream gating and analysis (**Supplemental Figure 1**).

### Analysis of tissue samples by immunofluorescent microscopy

*Sample collection:* Biopsies were collected using baby Tischler forceps at the lateral vaginal wall and the ectocervical os, embedded in optimal cutting temperature (OCT) compound and cryopreserved on dry ice.

*Laboratory Analysis*: Fresh frozen tissue biopsies in OCT compound were sectioned 8 μm thick using a cryostat and placed onto frosted microscope slides. Serial sections were used for H&E staining and immunofluorescent (IF) staining. H&E-stained slides were used to identify lamina propria and epithelium and to evaluate tissue integrity to ensure tissue sections met the criteria for IF analysis (intact lamina propria and epithelium).

Tissue sections for IF staining were fixed in acetone at -20°C for five minutes, dried for thirty minutes at room temperature, and rehydrated in tris-buffered saline with 0.05% tween (TBST). Slides were quenched using 3% H_2_O_2_ for twenty minutes at room temperature, rinsed with TBST, and endogenous binding sites were blocked using 1:10 fish gelatin blocking agent (Biotium, Cat# 22010) diluted in TBST for one hour. Unconjugated mouse anti-human CCR5 antibody (clone MC-5 (107) provided by M. Mack) at optimal dilution (0.3 µg/ml) was added for one hour, washed with TBST, and then goat anti-mouse antibody conjugated with horseradish peroxidase (polyclonal, Invitrogen, Cat# B40961) was added for thirty minutes. After washing with TBST, a solution of Tris Buffer (100mM, MilliporeSigma, Cat# 648315), optimally diluted tyramide (AF488, Invitrogen, Cat# B40953, diluted 1:100), and H_2_O_2_ (diluted 1:67,000) was then used for immunofluorescence and signal amplification. Slides were washed with TBST and a combination of optimally diluted, directly conjugated anti-CD3 (AF594, Clone UCHT1, R&D Systems, Cat# FAB100T, diluted 1:50) and anti-CD4 (AF647, Clone RPA-T4, BioLegend, Cat# 300520, diluted 1:25) antibodies were added for incubation overnight in the 4°C fridge. Slides were then rinsed in TBST and stained with DAPI (Invitrogen, Cat# D1306, diluted 2.5 ng/ml) for five minutes. Slides were washed with PBS, saturated briefly for ten seconds in ammonium acetate, saturated for ten minutes in copper sulfate, and then saturated again briefly for ten seconds in ammonium acetate. Slides were thoroughly rinsed in dH_2_O, dried, mounted with Prolong Gold (Invitrogen, Cat# P36930) and a cover slip, and allowed to set for 24 hours.

Tissue sections were imaged within 48 hours of staining using a 20x/0.8 Pan-APOCHROOMAT air objective on a Zeiss Axio Imager Z2 microscope as part of a TissueFAXS system (TissueGnostics; Vienna, Austria) equipped with an X-cite 120Q lamp (Excelitas), and with DAPI (49000 ET), EGFP (49002 ET), Texas Red (49008 ET), and Cy5 (490006 ET) filter sets. Images were acquired using an Orca-Flash 4.0 camera (Hamamatsu) with TissueFAXS 7.1 software (TissueGnostics).

### Tissue section image analysis

We quantified the density of CD3^+^, CD3^+^CD4^+^, and CD3^+^CD4^+^CCR5^+^ cells in the epithelium and lamina propria of CX and VT sections from individuals with or without BV. Images acquired using the TissueFAXS system were exported as multi-channel 16-bit tiff files of stitched regions. Cell segmentation, region of interest (ROI) selection (lamina propria vs epithelium), marker intensity thresholding, and quantification of densities were estimated using a custom graphical user interface (GUI) developed in MATLAB (R2022b). Nucleus segmentation was performed by finding the regional maxima of the grayscale DAPI signal, followed by morphological thickening constrained by the binary DAPI mask. Threshold intensities for the markers of interest were set manually through the GUI. The boundary between the lamina propria and epithelium regions was segmented based on nuclear density and further adjusted manually through the GUI. Finally, for each region (lamina propria and epithelium), the number of single-, double-, and triple-positive cells, as well as the density (number of cells per mm^2^), were extracted. Codes and an extended description of the GUI can be found at github.com/FredHutch/Kinga_Study_BV_MacLean.

### Cytokine and chemokine sample collection, and processing

*Cervicovaginal fluid:* The menstrual cup was inserted into the vaginal canal beneath the cervix at the beginning of the study visit and remained in place for a minimum of 15 and a maximum of 60 minutes while other visit procedures were being conducted. Sample collection was deferred if the participant was menstruating. The participant was encouraged to move around while the menstrual cup was in place. Once removed, the menstrual cup and all CVT fluid contents were placed in a 50 mL sterile collection tube and transported to the lab on ice. CVT fluids were collected in the 50 mL collection tube by centrifuging at 1500 rpm (320xg) for 10 min at 4°C. The mucous and fluids were gently mixed and 200-300 μl was aliquoted with a 1000 mL pipet tip into each cryovial and stored at -80°C.

*Serum*: Whole blood was collected in an SST vacutainer, allowed to clot for less than 2 hours, and centrifuged at 1300g for 15 minutes. Serum was aliquoted and stored at -80°C.

*Soluble mediator assays:* Serum and menstrual cup (CVT fluid) aliquots were shipped on dry ice to Eve Technologies (Calgary, Alberta, Canada). All samples were measured upon the first thaw. CVT fluid samples were weighed and diluted in PBS at a ratio of 1g sample to 1mL PBS. CVT fluid samples were vortexed for 2 minutes and up to 500 μl was loaded into 0.2 μm low-bind spin filters (MilliporeSigma, Cat# CWLS01S03) and centrifuged at ∼14,000 x g for 15 mins to remove the mucus from the samples prior to analysis. Levels of cytokines and chemokines from CVT fluid and serum samples were measured using the Human Cytokine Array/Chemokine Array 71-403 Plex Panel (Eve Technologies, HD71).

### Statistics

For purposes of data analysis, BV+ was defined by Nugent score of 7-10. BV- was defined as normal flora by Nugent score 0-3. For flow cytometry and soluble mediator analyses, to maximize the size of our BV+ group, we used enrollment visits of participants who were BV+ at enrollment, augmented with the six-month visit from any additional participants who were BV+ at the six-month exit visit. Thus, our BV+ group consisted of one time point per individual, but in some cases, it was enrollment while for others it was at exit. Among the remaining individuals, enrollment visits from those with normal flora at enrollment served as the BV- reference group. For immunofluorescent imaging, all specimens were from enrollment.

Cell densities of CD3^+^, CD3^+^CD4^+^, and CD3^+^CD4^+^CCR5^+^ cells in tissue sections were compared between individuals with versus without BV using the Wilcoxon rank sum test. To estimate adjusted differences in the percentage of T-cells with specified markers from flow cytometry data by BV status, we used rank-based regression, a nonparametric method robust to outliers (108). To investigate associations of soluble mediators in serum and CVT fluid samples with BV status, we first determined whether, for each mediator, at least 80% of the specimens generated a quantifiable level (vs out of quantifiable range or missing for another reason). If at least 80% of results were quantifiable, we imputed out of range values by randomly selecting a value between the lowest observed value and half the lowest observed value (if out of range low), or by selecting the largest observed value (if out of range high). We then compared log_10_- transformed levels of the mediator from individuals in BV+ vs BV- groups using a t-test and estimated differences in mean log cytokine concentrations from both serum and CVT fluid samples using linear regression. If <80% of the levels for a mediator were quantifiable, we categorized the mediator as detected vs not detected and estimated the odds ratio for the effect of BV on mediator detection using logistic regression. For both flow cytometry and soluble mediator data, results from VT and CX biopsy and CVT fluid samples were *a priori* adjusted for hormonal contraceptive use (yes, no and menstruating, no and amenorrheal), HSV-2 serology (positive, negative, indeterminate), HIV exposure (HIV status of sexual partner), and number of unprotected sex acts in the last thirty days (continuous), while results from PBMC samples were *a priori* adjusted for hormonal contraceptive use (yes, no and menstruating, no and amenorrheal), only. In this exploratory work, we did not adjust p-values for multiple testing. Statistical analyses were conducted using R version 4.3.3.

### Study approval

All participants provided written informed consent using documents reviewed and approved by the University of Washington institutional review board, and the Scientific and Ethics Review Unit of the Kenya Medical Research Institute.

## Data availability

All supporting data are included in the supplemental figures and tables and/or are available upon reasonable request to the senior authors.

## Author contributions

FM, JBG, JLS, SCV, NP, ICT, and LW conducted the experiments. FM, ATT, JLS, AS, and KKT analyzed data, MM provided reagents, JD and LKS contributed analysis methods, BHC, KN, NM, JRL, and JML designed the research study, and FM, MCS, JRL, and JML wrote the first draft of the manuscript. All authors edited and approved the manuscript.

## Acknowledgments

We thank all study volunteers for their participation in the Kinga Study and their willingness to provide many different samples. Additionally, we thank the members of the Kinga Study team and the Lund and Prlic labs for their helpful discussions on experimental findings and manuscript preparation throughout this process. Additionally, we would like to thank the Fred Hutchinson Cancer Center Cellular Imaging Shared Resource, supported by the Fred Hutchinson Cancer Center Cellular Imaging Core Facility (RRID:SCR_022609) of the Fred Hutch/University of Washington/Seattle Children’s Cancer Consortium (P30 CA015704), for assistance with microscopy and image analysis. This work was supported by the following grants from the National Institutes of Health: R01 AI131914 (to JML and JRL), R01 AI141435 (to JML), and R01 AI129715 (to JRL). SV was supported by T32 AI007509, ICT was supported by T32 AI007140 and LW was supported by T32 AI083203.

## Kinga Study Team

### University of Washington: International Clinical Research Center

Jairam R Lingappa (co-Principal Investigator and Protocol Chair), Justice Quame-Amaglo (study coordinator), Harald Haugen, Elena Rechkina, Daphne Hamilton, Matthew Ikuma, Marie Bauer, Xuanlin Hou, Zarna Marfatia, Katherine Thomas, Ayumi Saito, Adino Tesfahun Tsegaye

### Fred Hutchinson Cancer Center

Jennifer Lund (co-Principal Investigator), Jessica Graham, Jessica Swarts, Sarah Vick, Nicole Potchen, Irene Cruz Talevara, Lakshmi Warrier, Finn MacLean, Elizabeth McCarthy

### Kenya Medical Research Institute: Thika Partners in Health Research and Development (and Jomo Kenyatta University [JKUAT])

Nelly R. Mugo (Site Principal Investigator), Kenneth Ngure (Site Investigator), Catherine Kiptinness (site coordinator), Bhavna H. Chohan (Site Laboratory Director), Nina Akelo, Charlene Biwott, Stephen Gakuo, Elizabeth Irungu, Marion Kiguoya, Edith Kimani, Eric Koome, Solomon Maina, Linet Makena, Sarah Mbaire, Murugi Micheni, Peter Michira, Jacinta Nyokabi, Peter Mogere, Richard Momanyi, Edwin Mugo, Caroline Senoga, Mary Kibatha, Jelioth Muthoni, Euticus Mwangi, Philip Mwangi, Margaret Mwangi, Charles Mwangi, Stanley Mugambi Ndwiga, Peter Mwenda, Grace Ndung’u, Faith Njagi, Zakaria Njau, Irene Njeru, John Njoroge, Esther Njoroge, John Okumu, Lynda Oluoch, Judith Achieng Omungo.

---

**Supplementary Figure 1. Flow cytometry gating to determine phenotype expression on individual cells.**

Cells were gated by forward scatter height (FSC-H) and forward scatter area (FSC-A) to filter out non-singlet cells. Then cells were gated by side scatter area (SSC-A) and forward scatter area (FSC-A) to isolate lymphocytes. Lymphocytes were analyzed by live/dead and CD45 expression. Live CD45^+^ cells were then analyzed for CD3 expression. CD3^+^ cells were then analyzed for CD4 and CD8 expression. CD8^+^CD4^-^ were analyzed for the phenotypes listed. CD4^+^CD8^-^ were analyzed for CD25 and CD127 expression. CD25^-^ cells were defined as conventional CD4^+^ T cells (Tconv). CD25^+^CD127^low^ were analyzed for Foxp3 expression. Foxp3^+^ among the CD25^+^CD127^low^ population were defined as regulatory T cells (Tregs). Tregs were analyzed for the phenotypes listed. Tconv cells were analyzed for the phenotypes listed. In addition, Tconv cells were analyzed for T-bet expression which defined Th1 cells, and CCR6^+^CD161^+^ co-expression which defined Th17 cells. Th1 and Th17 cells were also analyzed for expression of the phenotypes listed. For each gating analysis, samples were only continued for further analysis if the parent gate had a minimum of 25 cells.

**Supplemental Figure 2. HIV target cell abundance was not altered by BV across CX and VT tissue compartments.**

Comparison of the cell density of CD3^+^ T cells, CD3^+^CD4^+^ T cells, or CD3^+^CD4^+^CD5^+^ HIV target cells between BV- and BV+ samples in the ectocervix lamina propria (**A**) and epithelium **(B)** and vaginal tract lamina propria (**C**) and epithelium (**D**). Wilcoxon rank sum test was performed for each comparison shown. Comparisons with p > 0.05 labeled “ns” for not significant. Each dot represents a measurement from an individual sample. Each horizontal bar indicates the median for its respective group. For each comparison, the Ns, medians, p-values (adjusted and unadjusted), and estimated differences (adjusted and unadjusted) are provided in Supplemental Table 2.

**Supplemental Figure 3. BV was associated with altered detection of soluble factors in CVT fluid and serum.**

Luminex was used to quantify cytokines and chemokines from CVT fluid or serum samples. Adjusted odds ratio (BV positive/BV negative) for CVT fluid cytokines (**A**) and serum (**B**) that did not meet the criteria for quantification of cytokine concentrations (fewer than 80% of samples were detectable). The adjusted 95% confidence interval is shown for all comparisons. Significant results when adjusted p ≤ 0.05 comparing BV- vs BV+ are colored in orange and non-significant differences (p > 0.05) comparing BV- vs BV+ are purple. Vertical dashed line at x=1 for reference. Serum comparisons were *a priori* adjusted for hormonal contraceptive use, and CVT fluid comparisons were *a priori* adjusted for hormonal contraceptive use, HSV-2 serology, HIV exposure, and semen exposure to reduce the effects of potential confounding variables on the analysis of BV-driven T cell alterations. Statistical analyses can be found in Supplemental Table 3.

## Table Legends

**Supplemental Table 4.** Antibodies used for high parameter flow cytometry on mucosal tissue biopsies and PBMCs.

