## Supplemental Figures 1-3 and Supplemental Table 4 for "Bacterial vaginosis-driven changes in cervicovaginal immunity that expand the immunological hypothesis for increased HIV susceptibility"

Supplemental Figure 1

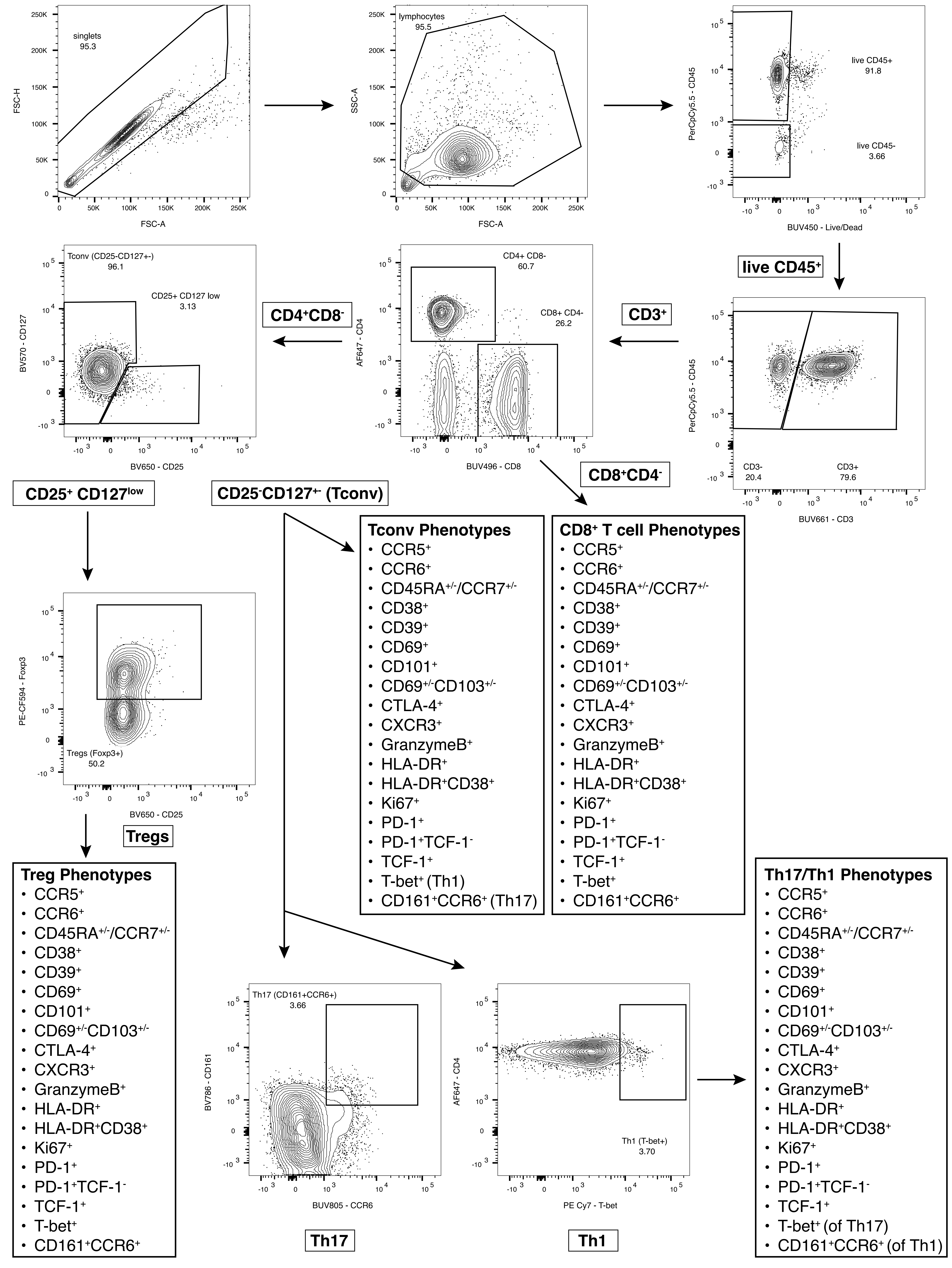

Supplemental Figure 2

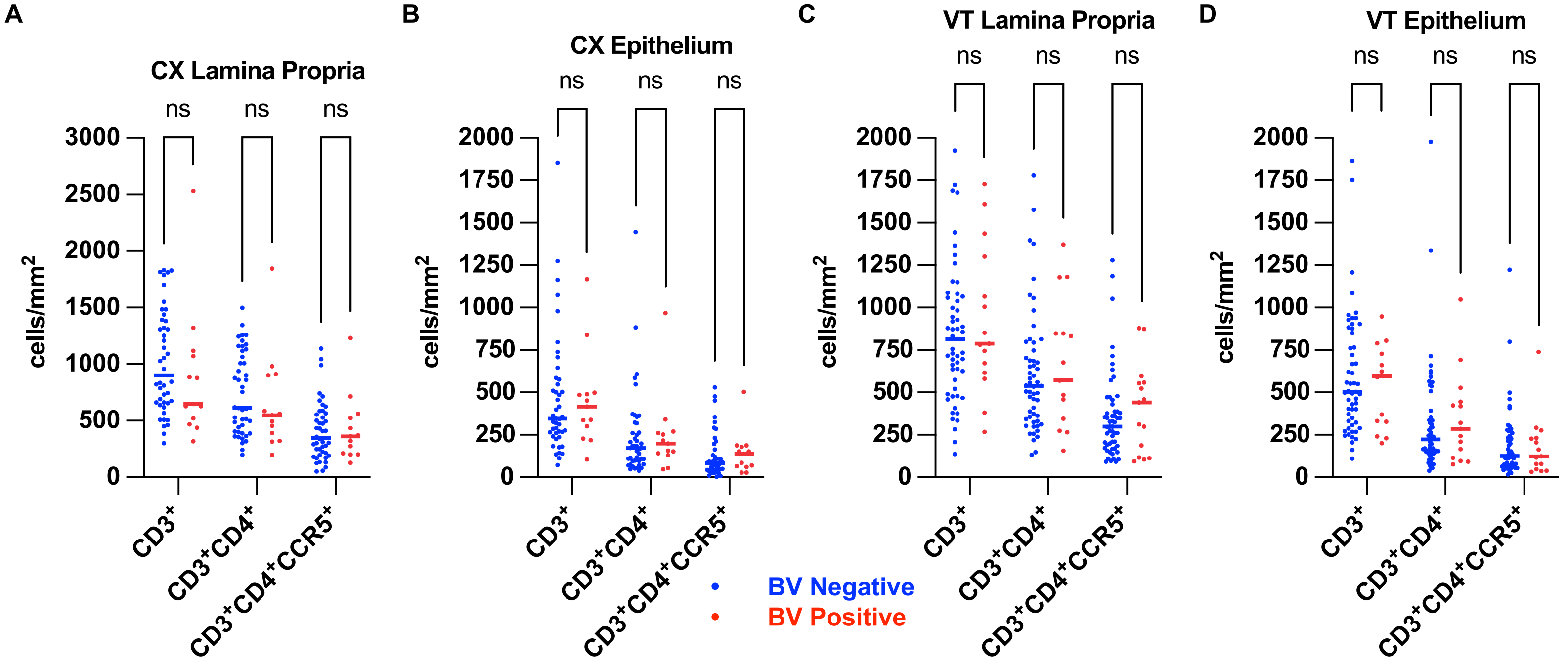

### Supplemental Figure 3

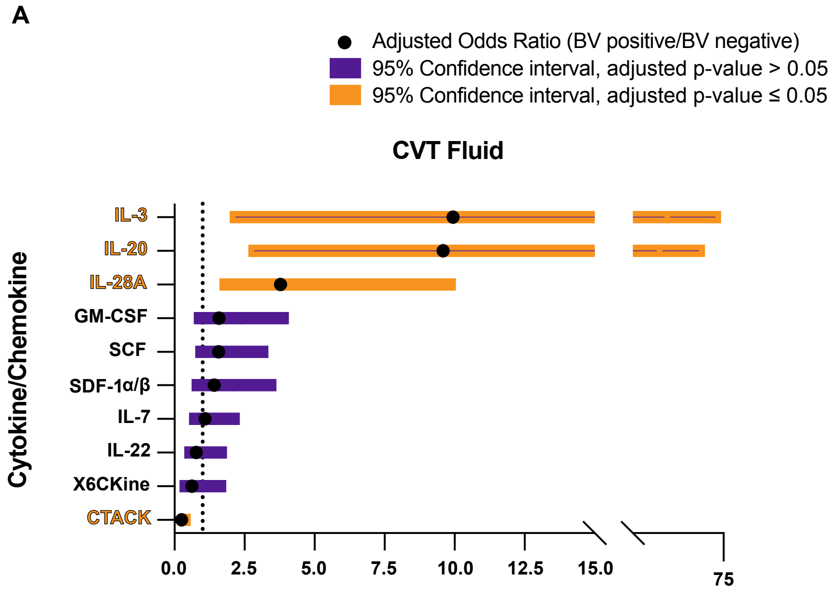


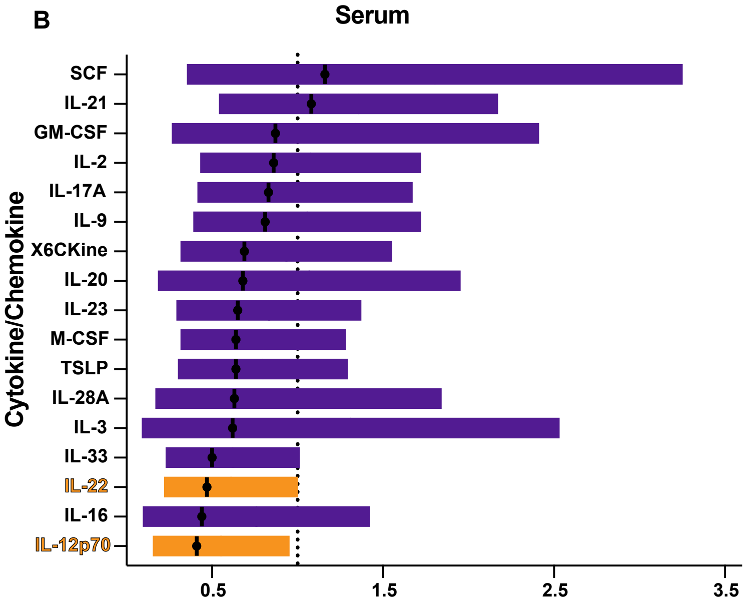

**Supplemental Table 4**

| T cell panel used for phenotyping | | | | | | |
|---|---|---|---|---|---|---|
| Channel | Fluorochrome | Antibody | Clone | Dilution | Company | Catalog # |
| UV395 | BUV395 | CD39 | TU66 | 1:40 | BD | 742524 |
| UV450 | BUV450 | Live/Dead | | 1:1000 | Invitrogen | L23105A |
| UV500 | BUV496 | CD8 | RPA-T8 | 1:500 | BD | 612943 |
| UV570 | BUV563 | CCR7 | 2-L1-A | 1:40 | BD | 749679 |
| UV660 | BUV661 | CD3 | UCHT1 | 1:200 | BD | 612964 |
| UV730 | BUV737 | CD69 | FN50 | 1:50 | BD | 612817 |
| UV780 | BUV805 | CCR6 | 11A9 | 1:20 | BD | 749361 |
| V510 | BV510 | CCR5 | J418F1 | 1:166.7 | BioLegend | 359128 |
| V570 | BV570 | CD127 | A019-D5 | 1:83.3 | BioLegend | 351308 |
| V610 | BV605 | PD-1 | EH12.2H7 | 1:20 | BioLegend | 329924 |
| V650 | BV650 | CD25 | BC96 | 1:83.3 | BioLegend | 302634 |
| V710 | BV711 | CD38 | HIT2 | 1:40 | BD | 563965 |
| V750 | BV750 | CD103 | Ber-ACT8 | 1:250 | BD | 747099 |
| V785 | BV786 | CD161 | HP-3G10 | 1:83.3 | BD | 748281 |
| B515 | AF488 | HLA-DR | 1.243 | 1:83.3 | BioLegend | 307656 |
| B660 | BB660 | SA | | 1:500 | BD | 6464295 |
| B710 | PerCpCy5.5 | CD45 | HI30 | 1:100 | BD | 564105 |
| B780 | BB790 | CD101 | V7.1 | 1:166.7 | BD | Custom |
| G575 | PE | TCF-1 | C63D9 | 1:40 | Cell Signaling | 14456 |
| G610 | PE-CF594 | Foxp3 | 236A/E7 | 1:20 | BD | 563955 |
| G660 | PE Cy5 | CTLA4 | BNI3 | 1:166.7 | BD | 561717 |
| G780 | PE Cy7 | T-bet | 4B10 | 1:20 | Invitrogen | 25-5825-82 |
| R660 | AF647 | CD4 | RPA-T4 | 1:40 | BD | 557707 |
| R710 | AF700 | GranzymeB | GB11 | 1:83.3 | BD | 560213 |
| R780 | APC-H7 | CD45RA | HI100 | 1:40 | BD | 560674 |
| B660 | Biotin | CXCR3 | G025H7 | 1:80 | BioLegend | 353743 |
